# Evolutionary and Functional Analysis of Monoamine Oxidase C (MAO C): A Novel Member of the MAO Gene Family

**DOI:** 10.1101/2024.08.20.608864

**Authors:** Gianluca Merello Oyarzún, Montserrat Olivares-Costa, Lorenzo Basile, Tammy P. Pástor, Pablo Mendoza-Soto, Luis Padilla-Santiago, Gonzalo A. Mardones, Claudia Binda, Juan C. Opazo

## Abstract

The monoamine oxidase (MAO) gene family encodes for enzymes that perform the oxidative deamination of monoamines, a process required to degrade norepinephrine, serotonin, dopamine, and other amines. While mammalian MAO enzymes, MAO A and MAO B, have been extensively studied, the molecular properties of the other family members are only partly uncovered. This study aims to explore the evolution of monoamine oxidases, emphasizing understanding the MAO gene repertoire among vertebrates. Our analyses show that the duplication that gave rise to MAO A and MAO B occurred in the ancestor of tetrapods, between 408 and 352 million years ago. Non-tetrapod jawed vertebrates possess the ancestral preduplicative condition of MAO A/B. Our results also identified a new family member, MAO C, in non-tetrapod jawed vertebrates. Thus, most jawed vertebrates possess a repertoire of two MAO genes, MAO A and MAO B in tetrapods and MAO A/B and MAO C in non-tetrapod jawed vertebrates, representing different MAO gene lineages. Jawless vertebrates possess the ancestral condition of a single copy gene, MAO A/B/C. Enzymatic assays conducted on the MAO recombinant enzymes of the Indo-Pacific tarpon show that both proteins, MAO A/B and MAO C, have enzymatic and molecular properties more similar to human MAO A, with the former featuring a strikingly higher activity rate when compared to all other MAO enzymes. Our analyses underscore the importance of scanning the tree of life for new gene lineages to understand phenotypic diversity and gain detailed insights into their function.

## Introduction

Today, the genomic field is in a golden age. The availability of genome sequences in representative species in all main branches of the tree of life represents a unique opportunity to study the genetic basis of phenotypic diversity. The release of hundreds of mammalian genomes by the Zoonomia project (Vignieri, 2023) and more than two hundred primate genomes represent just recent examples (Kuderna et al., 2023). With these data, we have learned, among other things, that variation in gene family size is widespread (Demuth et al., 2006), and a hidden gene diversity is waiting to be discovered (Himmel and Cox, 2020; Opazo et al., 2023).

In the brain, the mammalian monoaminergic system comprises neuromodulators, transporters, anabolic/catabolic enzymes, and receptors. Monoamine oxidases (MAOs) support the oxidative deamination of monoamines, a process necessary for the degradation of norepinephrine, serotonin, dopamine, and other amines. Two MAO proteins, MAO A and MAO B, inserted in the outer mitochondrial membrane have been described and classified according to their biochemical profile (Bach et al., 1988). While MAO A preferentially metabolizes serotonin, MAO B is selective towards phenylethylamine and benzylamine. Dopamine, epinephrine, norepinephrine, tryptamine, and tyramine are shared substrates for both enzymes (Shih et al., 1999).

MAO activities have been studied for decades using biological material from different animal species, including birds (Nicotra et al., 2002; Yeung et al., 2019), frogs (Kobayashi et al., 1981), fish (Anichtchik et al., 2006), and lampreys (Iagodina and Basova, 2013). In these studies and others, substrate specificity and inhibitor sensitivity profiles have allowed researchers to classify MAO proteins according to their resemblance to mammalian MAO A and B. Although informative, these studies have not considered a systematic analysis of MAO gene evolution. Attempts to reconstruct MAO phylogeny in metazoans have already been made (Anctil, 2009; Goulty et al., 2023; Kutchko and Siltberg-Liberles, 2013). Although these analyses included vertebrate MAOs, their sampling strategy was either designed to resolve phylogenetic relationships at a higher level (bilaterians or metazoans) or not comprehensive enough to reconstruct phylogenies in vertebrates with high confidence. Therefore, so far no study has focused exclusively on MAO evolution in vertebrates.

This study takes advantage of the golden age of genomics to advance our understanding of the evolution of MAOs in vertebrates. Our main results show that the duplication event that gave rise to MAO A and MAO B occurred in the ancestor of tetrapods. Non-tetrapod jawed vertebrates possess the ancestral preduplicative condition of MAO A/B. Interestingly, we identified a new monoamine oxidase gene family member in jawed vertebrates, MAO C. We experimentally show that, as expected, MAO A/B and MAO C localize to mitochondria. Enzymatic assays performed on the recombinantly produced MAOs of Indo-Pacific tarpon (IPT), as an example of a species possessing the MAO A/B and MAO C, show that both IPT-MAO A/B and IPT-MAO C have enzymatic and molecular properties more similar to human MAO A, with the former featuring a strikingly higher activity rate with respect to all MAO enzymes. Finally, bioinformatic analyses of brain expression patterns in the sea lamprey indicate that the vertebrate ancestral MAO protein probably was involved in vascular cell and monoaminergic neuron function.

## Results and Discussion

### Monoamine oxidase A and B genes originated in the ancestor of tetrapods

We performed phylogenetic analyses to investigate the evolutionary origin of MAO A and MAO B genes. Our maximum-likelihood tree recovered a well-supported clade containing MAO A sequences from tetrapods (Fig. 1, blue shading). Synteny analyses on the locus containing MAO A support our phylogenetic results (Supplementary Figure S1). Sequences recovered in the MAO A clade correspond to the MAO gene located at the 3’ side of the GPR82 gene (Supplementary Figure S1). Interestingly, MAO A underwent a duplication event in the last common ancestor of anurans, giving rise to two MAO A genes (Fig. 1).

**Figure 1.**
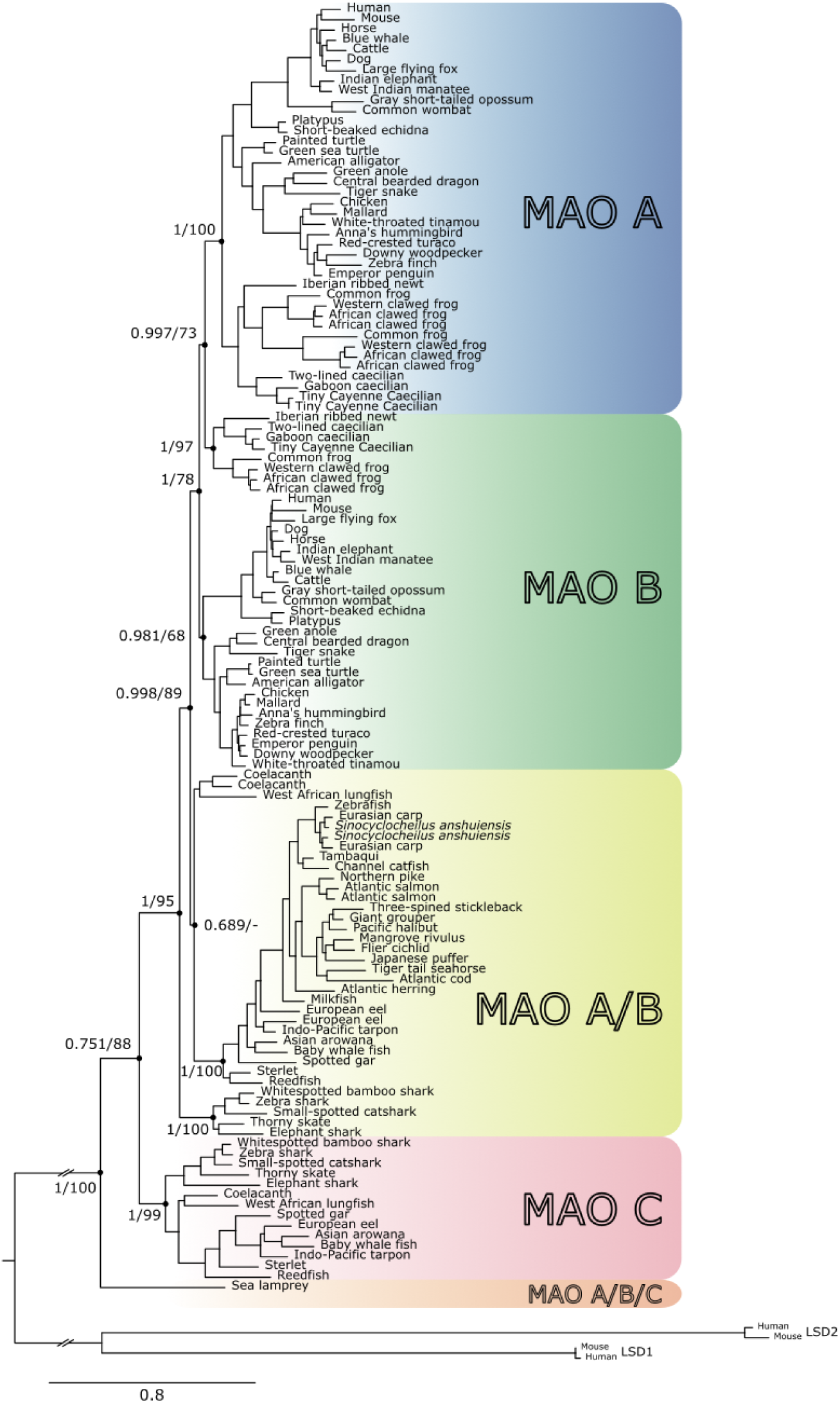
Maximum-likelihood tree showing relationships among monoamine oxidases of vertebrates. Numbers on the nodes correspond to support from the abayes and ultrafast bootstrap values. The scale denotes substitutions per site, and shading represents MAO lineages. Lysine-specific demethylase 1A and 1B of the human (*Homo sapiens*) and house mouse (*Mus musculus*) were used as outgroups.

MAO B sequences were not recovered monophyletic (Fig. 1, green shading). We recovered a clade containing MAO B sequences from mammals, birds, and reptiles and another clade containing MAO B sequences from amphibians (Fig. 1). The clades we identify as MAO B correspond to the MAO gene located at the 5’ side of the NDP syntenic gene, and at the 3’ of MAO A (Supplementary Figure S1). The lack of monophyly could be explained by the difficulty of resolving phylogenetic relationships with a small number of amino acid changes, as revealed by the shortness of some branches in our phylogenetic tree (Fig. 1). A recent study also had difficulty resolving the MAO B monophyly (Goulty et al., 2023).

We performed additional phylogenetic analyses using nucleotide sequences to further explore the monophyly of MAO B (Supplementary Figure S2). In these new analyses, we recover the monophyly of both paralogs in tetrapods (Supplementary Figure S2), supporting the idea that the lack of monophyly in the amino acid tree could be due to the lack of information. Thus, by combining information from our synteny and phylogenetic analyses, we could recognize MAO A and MAO B lineages only in representative species of tetrapods (Fig. 1 and Supplementary Figures S1 and S2). In our survey, we noticed that MAO genes are mainly located on autosomes, while in humans and other mammals, they are located on chromosome X, as previously noticed (Pintar et al., 1981). To shed light on when the translocation occurred, we retrieved the chromosomal location in species representative of all main groups of tetrapods. According to our results, MAO genes were translocated to chromosome X in the ancestor of placental mammals between 160 and 99 million years ago (Kumar et al., 2022) (Supplementary Figure S3).

In non-tetrapods, we recovered a clade containing MAO sequences from the West African lungfish, coelacanth, and bony fish sister to the clade containing MAO A and MAO B from tetrapods (Fig. 1). Cartilaginous fish MAO sequences were sister to all other sequences (Fig. 1). We named these clades MAO A/B (Fig. 1, yellow shading). In almost all non-tetrapod species, we found a single copy gene (MAO A/B) flanked by the same genes as MAO A and MAO B in tetrapods (Supplementary Figure S1), suggesting that the duplication event that gave rise to MAO A and MAO B genes occurred in the last common ancestor of tetrapods between 408 and 352 million years ago (Kumar et al., 2022).

### Identification of a new Monoamine Oxidase Gene Family Member, MAO C, in non-tetrapod jawed vertebrates

Unexpectedly, our analysis identified a new MAO gene family member, MAO C (Fig. 1, pink clade). This new MAO paralog was found in the genome of the West African lungfish, coelacanth, bony fish, and cartilaginous fish (Fig. 1, pink clade), and its monophyly is strongly supported (1/99; Fig. 1, pink clade). The MAO C clade was recovered sister to the clade containing MAO A, MAO B, and MAO A/B sequences from jawed vertebrates (Fig. 1). Synteny analyses reveal that MAO C present in bony fish is flanked at the 5’ side by the CMC4 and BRCC3 genes, and by the GABRA3 and GABRB4 genes at the 3’ side (Supplementary Figure S1). In the case of cartilaginous fish, synteny is maintained at the 5’ side and partially conserved at the 3’ side (Supplementary Figure S1). MAO C in the West African lungfish and coelacanth does not show conserved synteny. However, they do share some genes that are neighboring to each other (Supplementary Figure S1). In support of our results, a recent study of the evolution of 18 genes involved in the synthesis, turnover, and detection of monoamines in animals recovered a clade of two MAO sequences from jawed vertebrates, sister to the jawed vertebrate clade containing MAO A and MAO B, insinuating the presence of a new MAO gene lineage in vertebrates (Goulty et al., 2023). Finally, the sea lamprey (*Petromyzon marinus*) sequence was recovered sister to all jawed vertebrate sequences previously mentioned (we named this family member MAO A/B/C) (Fig. 1).

Our model for the evolution of the monoamine oxidases in vertebrates is graphically summarized in Figure 2. According to our model, the vertebrate ancestor had a single copy gene, an ancestral condition that is maintained in cyclostomes (Fig. 2). In the last common ancestor of jawed vertebrates, a duplication event gave rise to MAO A/B and MAO C gene lineages (Fig. 2). The MAO A/B and MAO C genes are present in representative species of cartilaginous fish, bony fish, coelacanth and West African lungfish (Fig. 2). MAO C was lost in the ancestor of tetrapods (Fig. 2), while MAO A/B duplicated giving rise to MAO A and MAO B (Fig. 2).

**Figure 2.**
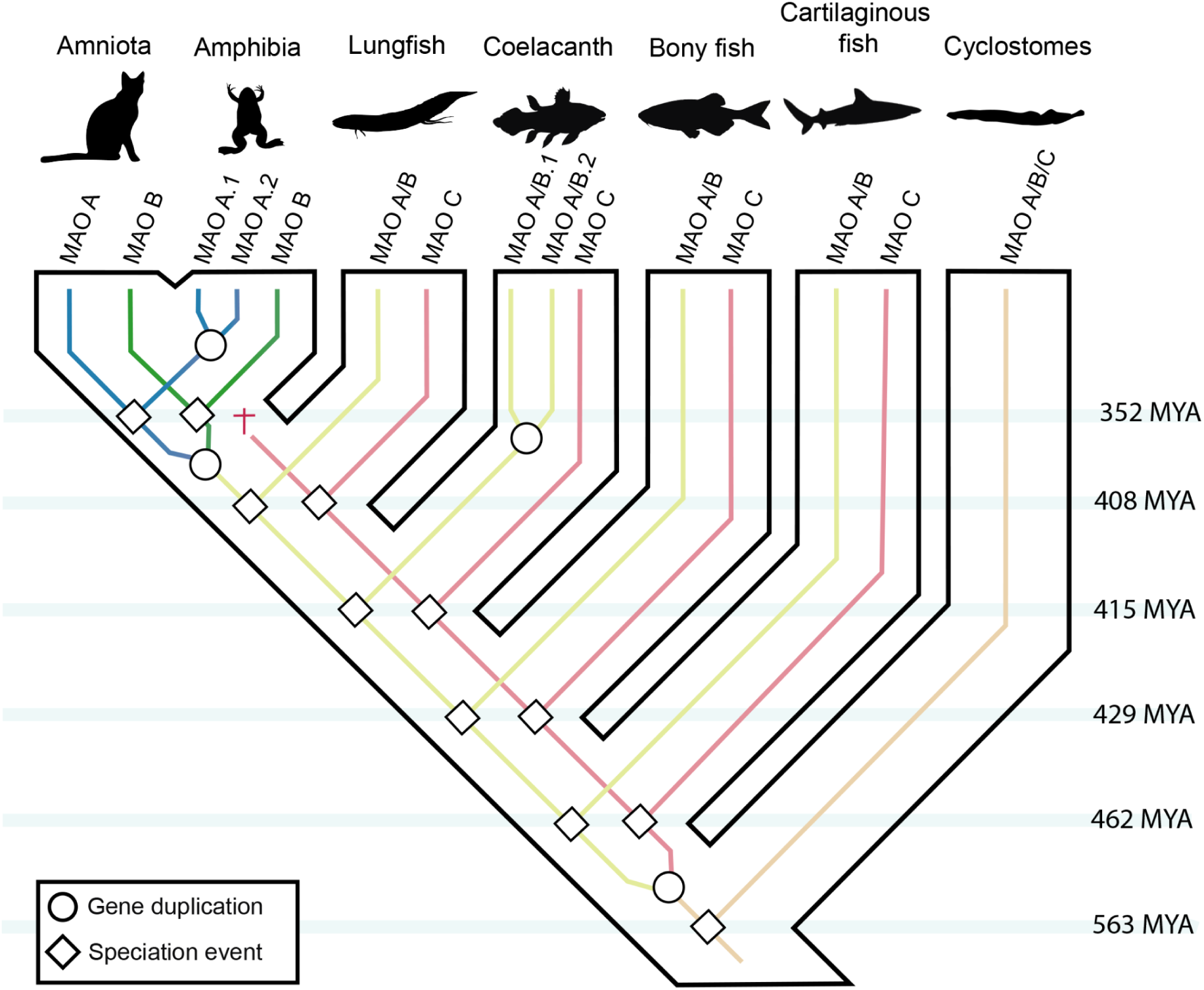
An evolutionary model regarding the evolution of the monoamine oxidases in vertebrates. Our model proposes that the vertebrate ancestor had a single copy gene (MAO A/B/C), a condition that cyclostomes maintain. This gene underwent a duplication event in the ancestor of jawed vertebrates, giving rise to MAO A/B and MAO C. These genes are found in cartilaginous fish, ray-finned fish, coelacanths, and West African lungfish. In the ancestor of tetrapods, the MAO C gene was lost. In contrast, the MAO A/B gene underwent a duplication event in the tetrapod ancestor, originating MAO A and MAO B. Silhouette images were downloaded from PhyloPic (http://phylopic.org/).

The description of new gene family members is not uncommon (Himmel et al., 2020; Opazo et al., 2021; Wichmann et al., 2016), especially in the present, when high-quality genomes of non-model species are being sequenced and deposited in public databases (Kuderna et al., 2023; Martin et al., 2023; Sayers et al., 2022; Vignieri, 2023). The high-throughput methodologies for inferring homology relationships have contributed to understanding the evolutionary dynamics of gene families (Altenhoff et al., 2024; Emms and Kelly, 2019). It has also contributed the manual curation by reconciling gene trees with species trees (Goodman et al., 1979). This way, the scientific community has realized that genetic variation, expressed as gene copy number variation, is widespread and represents an opportunity to gain insights into the genetic bases of phenotypic diversity (Chen et al., 2013). The case of the MAO genes is interesting as most jawed vertebrates possess the same number of genes, two MAO genes, but representing different MAO gene lineages (Fig. 2). Mammalian MAO A and MAO B are the most extensively studied members of the family (Edmondson and Binda, 2018; Jones and Raghanti, 2021; Shih et al., 1999). Beyond the brain, they are expressed at different levels throughout the body, with MAO A being unique in the placenta, while in platelets only MAO B is expressed (Edmondson and Binda, 2018). In agreement with a pattern of partial subfunctionalization after gene duplication, they deaminate neuroactive aromatic amines with a partly overlapping substrate specificity, including neurotransmitters and exogenous amines. These enzymes feature different inhibitor selectivity, an important aspect of their role as validated drug targets for neurological diseases (Edmondson and Binda, 2018). Much less is known for species that possess the other repertoire of two MAO genes. Functional characterization of species containing MAO A/B and MAO C will shed light on the consequences of gene repertoires with different evolutionary origins.

### MAO A/B and MAO C proteins localize to mitochondria

Mammalian MAOs are mitochondrial outer membrane enzymes that catalyze the oxidative deamination of substrates through a FAD-dependent mechanism ((Edmondson et al., 2009; Kearney et al., 1971; Wu et al., 1993). To determine whether MAO A/B and MAO C localization are conserved, we analyzed both proteins from the Indo-Pacific tarpon (*Megalops cyprinoides*). We expressed in human cultured cells the full-length proteins tagged with three copies of the c-Myc epitope at the N-terminus (3myc-MAO A/B and 3myc-MAO C). Fluorescence microscopy analysis of HeLa cells expressing either protein showed that the monoclonal antibody 9E10 against the c-Myc epitope labeled structures highly reminiscent of mitochondria (Fig. 3A and B). The mitochondrial localization of 3myc-MAO A/B and 3myc-MAO C was confirmed by their co-localization with the fluorescent signal from MitoTracker Orange (Pearson’s correlation coefficient [*r*] = 0.97 ± 0.03; *n* = 20) (Fig. 3A and B). To obtain biochemical evidence of the expression of each protein, we performed immunoblot analysis with monoclonal antibody 9E10, which showed the detection of expected 64 and 64.5 kDa proteins for MAO A/B or MAO C, respectively, in all cell lines analyzed (Fig. 3C). These results indicate that ectopically expressed MAO A/B and MAO C proteins of the expected size are targeted to the mitochondria.

**Figure 3.**
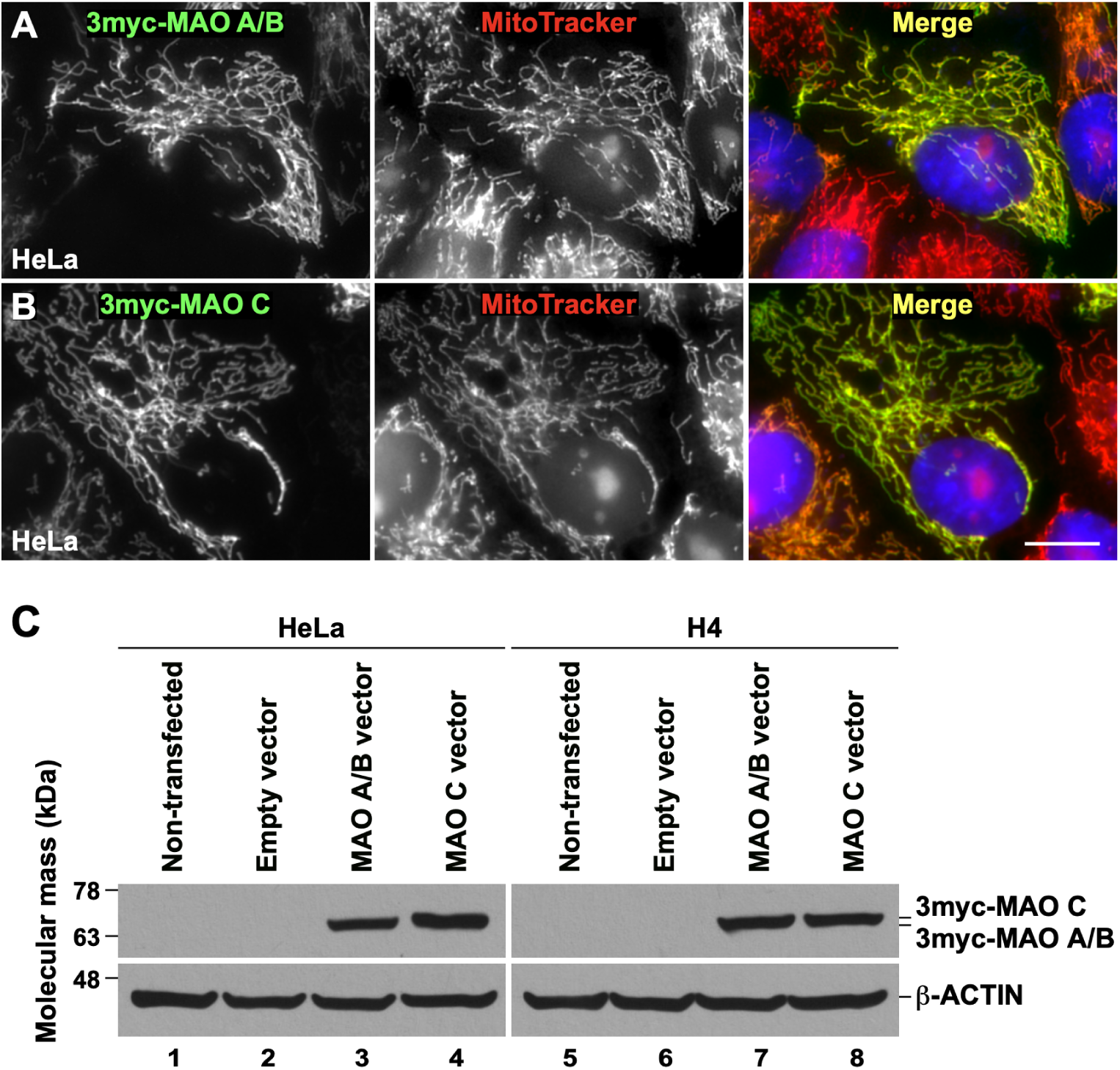
Expression of Indo-Pacific tarpon (*Megalops cyprinoides*) MAO A/B and MAO C in human cells. (A and B) HeLa cells grown in glass coverslips were transfected to express full-length MAO A/B (A) or MAO C (B) tagged with three copies of the c-Myc epitope at the N-terminus (3myc-MAO A/B and 3myc-MAO C, respectively), followed by incubation with the mitochondrial probe MitoTracker Orange (red channel). Cells were fixed, permeabilized, and labeled with mouse monoclonal antibody against the c-Myc epitope, followed by incubation with Alexa-Fluor-488-conjugated donkey anti-mouse IgG (green channels), and nuclei were stained with the DNA probe DAPI. Stained cells were examined by fluorescence microscopy. In the merge channels, yellow indicates the colocalization of each MAO protein with MitoTracker at mitochondria, and red depicts mitochondria in non-transfected cells. Bar, 10 μm. (C) The indicated cells were left untreated (Non-transfected; lanes 1 and 5), transfected with empty pCMV-3Tag-2A vector (lanes 2 and 6), or transfected with pCMV-3Tag-2A vector encoding either 3myc-MAO A/B (lanes 3 and 7) or 3myc-MAO C (lanes 4 and 8). Detergent-soluble extracts were prepared, and samples were processed by SDS-PAGE and immunoblot analysis with antibodies against the c-Myc epitope or antibody against β-ACTIN used as loading control. The position of molecular mass markers is indicated on the left.

### Enzymatic and molecular properties of Indo-Pacific tarpon MAO A/B and MAO C

While mammalian MAO enzymes have been thoroughly investigated, the molecular properties of the other family members in other vertebrates are only partly uncovered. A single gene encoding for a MAO enzyme was identified in zebrafish (*Danio rerio*) (Setini et al., 2005), which proved to be a reliable animal model system to identify neuroactive small molecules (Kokel et al., 2010). The biochemical characterization of zebrafish MAO showed that the enzyme displays a dual functionality, although its properties are more similar to human MAO A than to MAO B (Aldeco et al., 2011). The discovery of a new MAO gene family member in non-tetrapod jawed vertebrates in addition to MAO A and MAO B, prompted us to produce the two protein members existing in Indo-Pacific tarpon in their recombinant forms to study their enzymatic properties. Sequence analysis showed that both MAO A/B and MAO C from Indo-Pacific tarpon (IPT-MAO A/B and IPT-MAO C) have an overall sequence identity in the range of 63-72% in comparison to human MAOs, including the C-terminal 30 amino acids that form the outer mitochondrial membrane-spanning-helix. This observation is supported by the cell culture analysis described in the previous section, which showed that MAO A/B and MAO C localize in mitochondria (Fig. 3). By following similar procedures developed for the human MAOs (see Materials and Methods section), we produced recombinant Indo-Pacific tarpon MAOs using *Pichia pastoris* as heterologous expression system and purified the proteins as detergent-solubilized preparations.

Both IPT-MAO A/B and IPT-MAO C could be expressed and purified to homogeneity, yielding bright yellow samples, which indicates the presence of the FAD cofactor associated with the protein. In general, the reaction catalyzed by MAO enzymes consists of FAD-dependent oxidation of the C-N bond of primary or secondary (or tertiary in some cases) monoamine substrates, leading to an imine intermediate that is non-enzymatically hydrolyzed into the final aldehyde product (Edmondson et al., 2009). In each enzyme cycle, the reduced FAD cofactor is re-oxidized by molecular dioxygen, generating hydrogen peroxide as a secondary product. We performed a screening of the IPT-MAO enzymatic activity using different amine substrates, and for those that proved to be active, we determined the steady-state kinetic parameters (Table 1). In parallel, to enable comparative analysis, we conducted the same experiments with recombinant human MAO A and MAO B, as data on the various substrates were partially reported in the literature, and some inconsistencies might arise depending on different enzyme preparations. Both IPT-MAO A/B and IPT-MAO C are active on all tested substrates (Table 1). Interestingly, IPT-MAO C shows activity *k*_cat_ values comparable to those measured for the human enzymes, whereas IPT-MAO A/B is strikingly much more active on all substrates. Except for serotonin, this enzyme displays *k*_cat_ values that are about one order of magnitude higher, and in some cases this result is even more pronounced if taken as *k*_cat_/*K*_m_ ratio. In physiological conditions, substrate specificity of human MAOs depends on their expression pattern in different tissues. Nevertheless, serotonin and phenethylamine are considered specific substrates of MAO A and MAO B, respectively, which can be observed also when measuring enzymatic activity in buffer mixtures on purified proteins (Table 1). Our assays on IPT-MAOs show that both serotonin and phenethylamine are good substrates for either enzyme, provided the overall higher activity featured by IPT-MAO A/B as mentioned above. Although a clear “MAO A-like” or a “MAO B-like” behavior cannot be identified in any of the two IPT-MAOs, they are barely active on benzylamine that is a non-physiological substrate specifically oxidized by human MAO B (with very low activity with human MAO A). This observation is in agreement with the tests performed with the isoform-selective inhibitors clorgyline and deprenyl. Both compounds contain a propargylic moiety that promptly reacts with the FAD cofactor forming a covalent adduct that irreversibly inactivates the enzyme (De Colibus et al., 2005), being human MAO A and MAO B selectively inhibited by clorgyline and deprenyl, respectively. The formation of the covalent adduct can be monitored spectrophotometrically by following the disappearance of the typical 370 and 456 nm absorbance peaks of the oxidized flavin and the formation of a peak at 420 nm. Interestingly, both IPT-MAO A/B and IPT-MAO C react with clorgyline, with the latter being slightly slower in reaching the fully inactivated state (Figure 4). Deprenyl inactivates IPT-MAO A/B similarly to clorgyline, whereas IPT-MAO C is totally unaffected. These data agree with the observation that IPT-MAO A/B is highly reactive and suggest that this enzyme shares catalytic features with both human MAOs. Instead, IPT-MAO C displays a lower reactivity and is enzymatically more similar to human MAO A than to MAO B.

**Figure 4.**
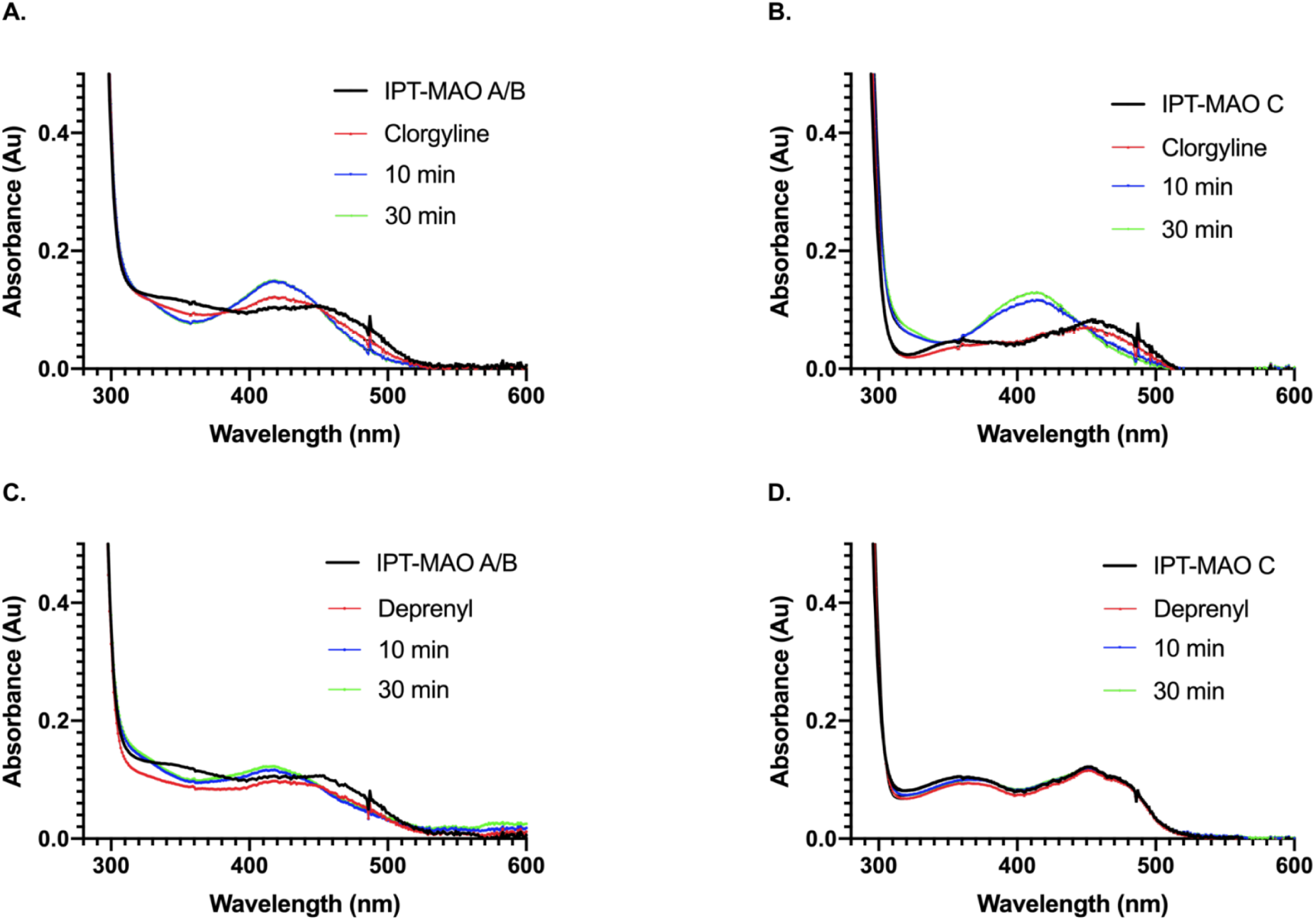
UV-Vis absorbance spectra of IPT-MAO A/B and IPT-MAO C (10 µM) incubated with 10-fold molar excess of either clorgyline or deprenyl. Measurements were registered on the native enzyme (black) immediately after the addition of the inhibitor (red) and at 10 (blue) and 30 (green) minutes.

**Table 1.**
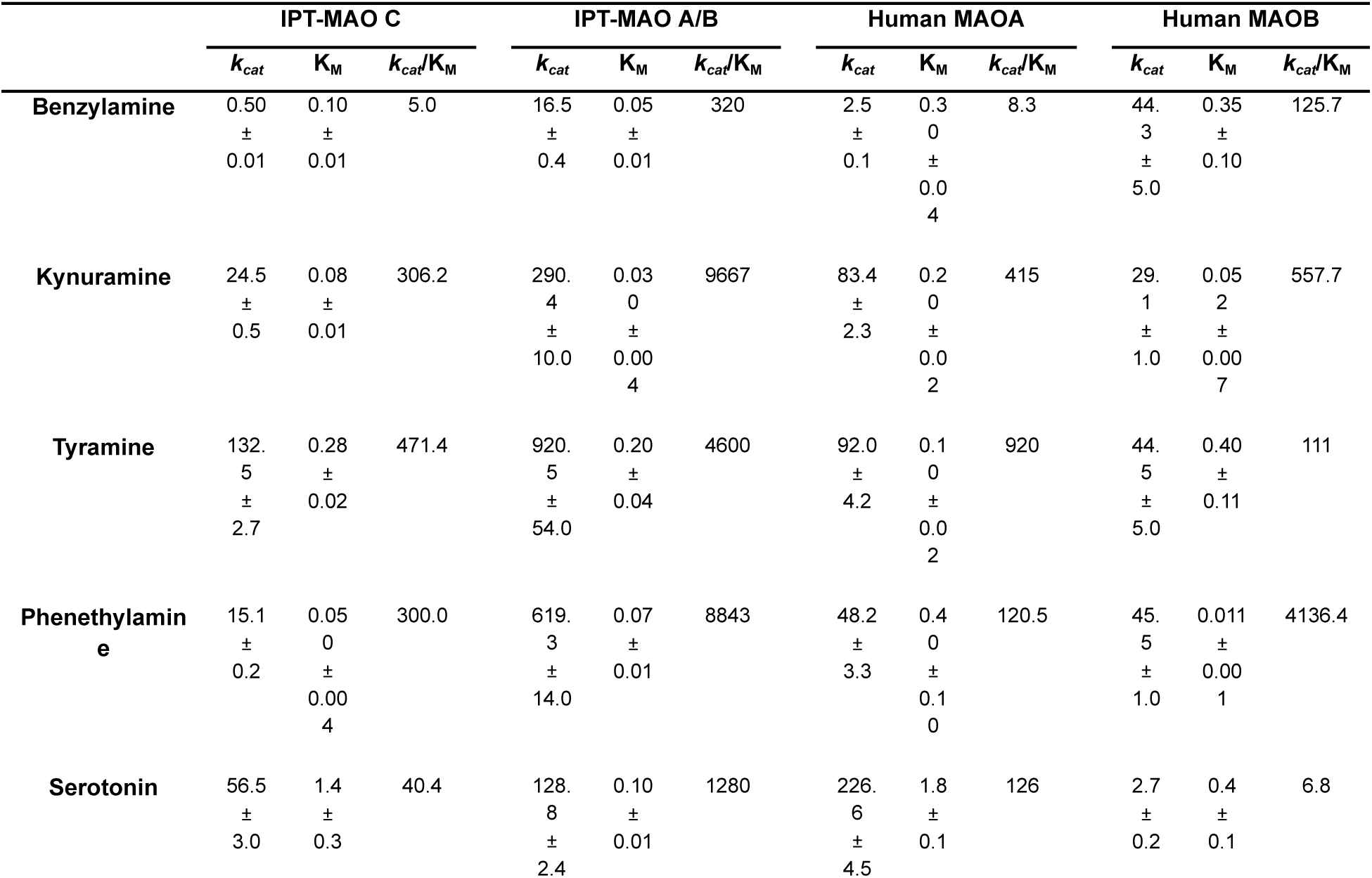
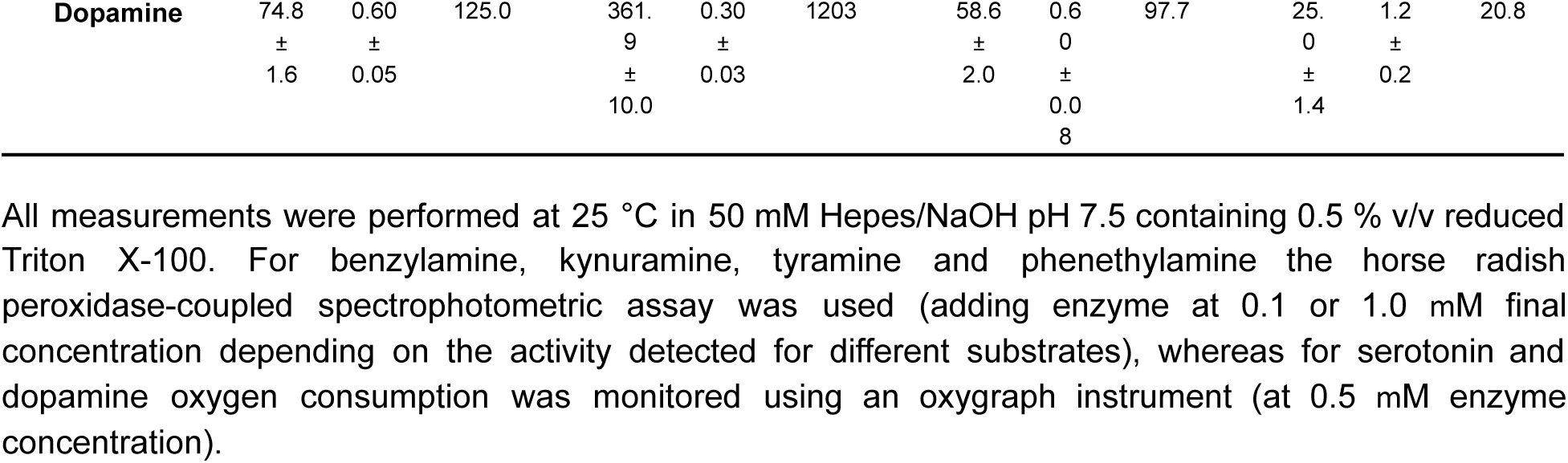
Kinetic parameters measured for the amine substrates with MAO enzymes. Values are expressed as min^-1^ and mM for *k_cat_* and K_M_, respectively.

All measurements were performed at 25 °C in 50 mM Hepes/NaOH pH 7.5 containing 0.5 % v/v reduced Triton X-100. For benzylamine, kynuramine, tyramine and phenethylamine the horse radish peroxidase-coupled spectrophotometric assay was used (adding enzyme at 0.1 or 1.0 mM final concentration depending on the activity detected for different substrates), whereas for serotonin and dopamine oxygen consumption was monitored using an oxygraph instrument (at 0.5 mM enzyme concentration).

To rationalize the kinetic data, we generated the 3D structure models of IPT-MAO A/B and IPT-MAO C by using AlphaFold3 (Abramson et al., 2024), which can accurately predict protein structures including complexes with cofactors. In both cases, the whole model scored a very high confidence value (pLDDT>90) except for the transmembrane C-terminal segment (90>pLDDT>70), which nevertheless is predicted to fold into an ɑ-helix as found in the crystal structures of human MAOs. Inspection of the active site structure of the two IPT enzymes superposed onto human MAO A and MAO B (Fig. 5) revealed that the area around the flavin site where substrate oxidation occurs is highly conserved. In particular, the Tyr-Tyr pair that forms the aromatic cage with the flavin ring and the position of FAD itself (in proximity of the Cys residue covalently linked to the cofactor in the crystal structures of human MAOs) are the same in all proteins, being this molecular architecture essential to accomodate the aromatic substrate in the correct way to allow the catalytic reaction. In addition, also the Lys residue on the edge of the flavin which is involved in dioxygen binding (Iacovino et al., 2020) adopts the same conformation, as well as the Tyr and Phe residues at the bottom and top of the catalytic niche. Differences arise in the area of the active site more distant from FAD, which also determines the divergent inhibitor selectivity between the human enzymes. The gating residues that in human MAO A are Phe208/Ile335 and in human MAO B are Ile199/Tyr326 correspond to a Phe/Leu in IPT-MAO A/B and Phe/Met in IPT-MAO C (Milczek et al., 2011). Moreover, Cys172 in human MAO B (known to form hydrogen bonds with selective inhibitors) corresponds to a Gln in both IPT MAOs like in human MAO A. Altogether, these observations indicate that all MAO hallmarks are present in both IPT enzymes and that their active site is more similar to human MAO A than to human MAO B, which is in agreement with the kinetic data. IPT-MAO C appears to have some unique features, such as a high content of Met residues throughout the sequence, including two in the active site. This is consistent with a separate gene family member whose role will be the subject of further investigations.

**Figure 5.**
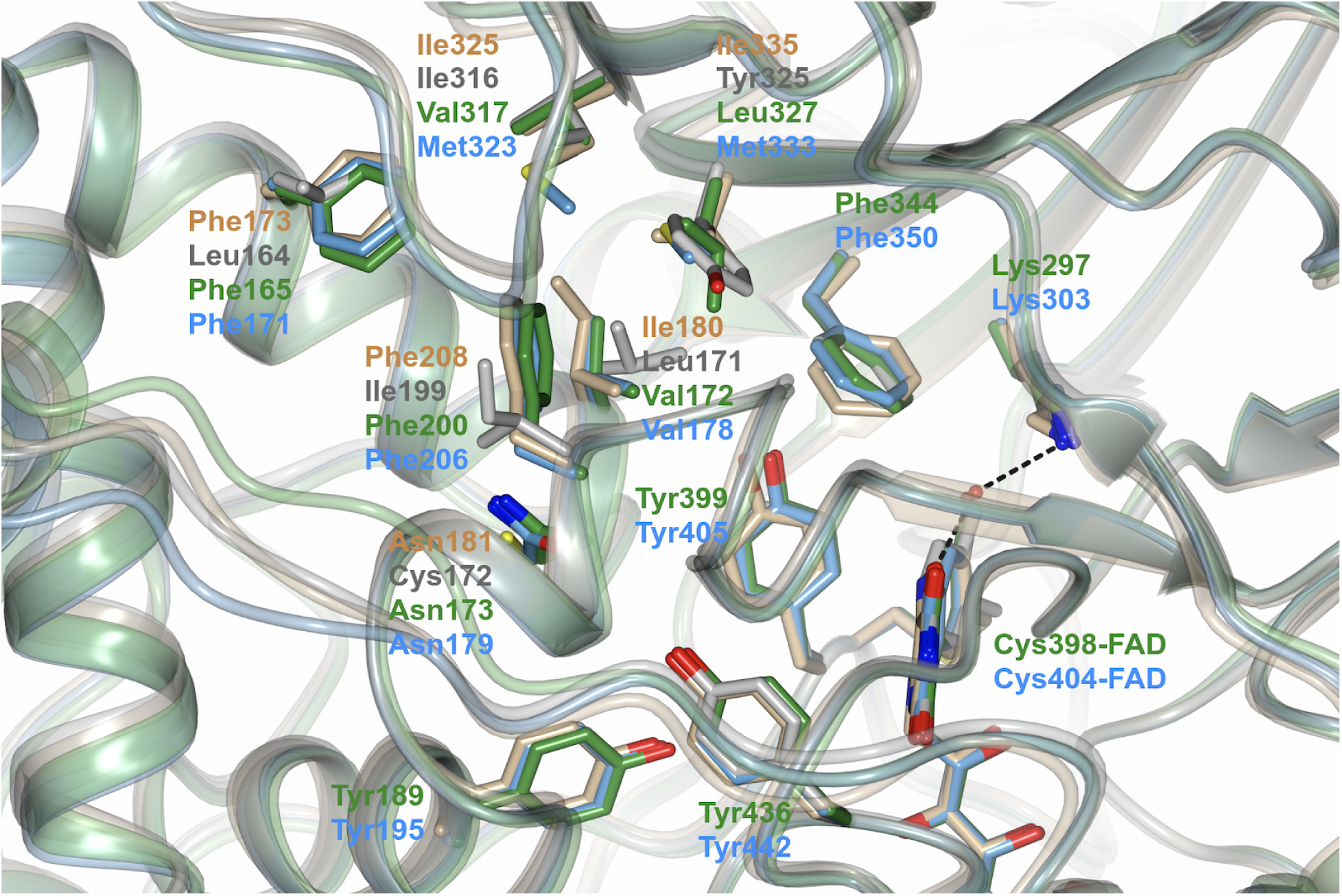
Zoomed view of the superposed 3D structures of MAO enzymes showing the active site. IPT-MAO A/B (green) and IPT-MAO C (light blue) are models predicted by AlphaFold3, whereas human MAO A (beige) and MAO B (gray) are crystal structures (PDB codes 2Z5X and 2V5Z, respectively; inhibitors bound to the structures were removed from the coordinate files). Structures are represented as semi-transparent ribbon diagram with FAD and the residues facing the active site represented as sticks (oxygen, nitrogen and sulfur atoms are red, blue and yellow, respectively). Labels are indicated according to the ribbon colors mentioned above; for residues that are fully conserved in all four proteins labels are reported only for the IPT homologues. In the crystal structures the water molecule bridging the conserved lysine residue (Lys305 in human MAO A and Lys296 in human MAO B) and the N5 atom of the FAD cofactor is shown as red sphere. The figure was prepared using the software CCP4mg (McNicholas et al., 2011).

### Brain expression of MAO genes in species with contrasting gene repertoires

Single-cell RNA sequencing (scRNA-seq) has enabled the classification of cell types based on their transcriptomic profile. Brain atlases obtained by scRNA-seq have allowed researchers to advance our understanding of brain evolution by comparing cell-type diversity in specific anatomical structures among different species. This resource is invaluable for understanding the expression patterns of different gene repertoires with single-cell resolution. In our work, we took advantage of the atlases of two species (house mouse, *Mus musculus* and sea lamprey, *Petromyzon marinus*) (Lamanna et al., 2023; Yao et al., 2023) in which all major divisions of the brain have been sequenced – telencephalon, diencephalon, mesencephalon, and rhombencephalon – to compare the cell types in which the ancestral condition of a single copy gene present in the sea lamprey (MAO A/B/C) and the duplicated genes in the house mouse (MAO A and MAO B) are expressed.

Our analysis showed widespread expression of MAO A/B/C in the sea lamprey brain (Fig. 6A). The highest abundance was observed in cell clusters identified as monoaminergic neurons, peptidergic neurons, and, especially, vascular cells. Clusters comprised of excitatory and inhibitory neurons of the telencephalon, habenular neurons, excitatory neurons of the diencephalon and mesencephalon, inhibitory neurons from the mesencephalon, spinal cord and rhomboencephalic neurons, and immune cells show moderate expression of MAO A/B/C. The choroid plexus epithelium, meningeal fibroblasts, and erythrocytes have no detectable expression of MAO A/B/C (Fig. 6A and B). In the case of the house mouse, MAO A is also widely expressed throughout the brain (Fig. 6C, left panel). Intriguingly, like in the sea lamprey, MAO A expression is highest in vascular cells. High expression was also observed in astrocytes, oligodendrocytes and oligodendrocyte precursor cells (OPC), habenular (HB) glutamatergic neurons, histaminergic neurons, serotonergic neurons and dopamine-β-hydroxylase (Dbh)-expressing neurons from the brainstem, among others (Fig. 6C left, Supplementary Figure S4). MAO B expression is more restricted. The highest expression is observed in serotonergic neurons, histaminergic neurons, thalamic (TH) glutamatergic neurons, vascular cells, and astrocytes (Fig. 6C right panel, Supplementary Figure S4).

**Figure 6.**
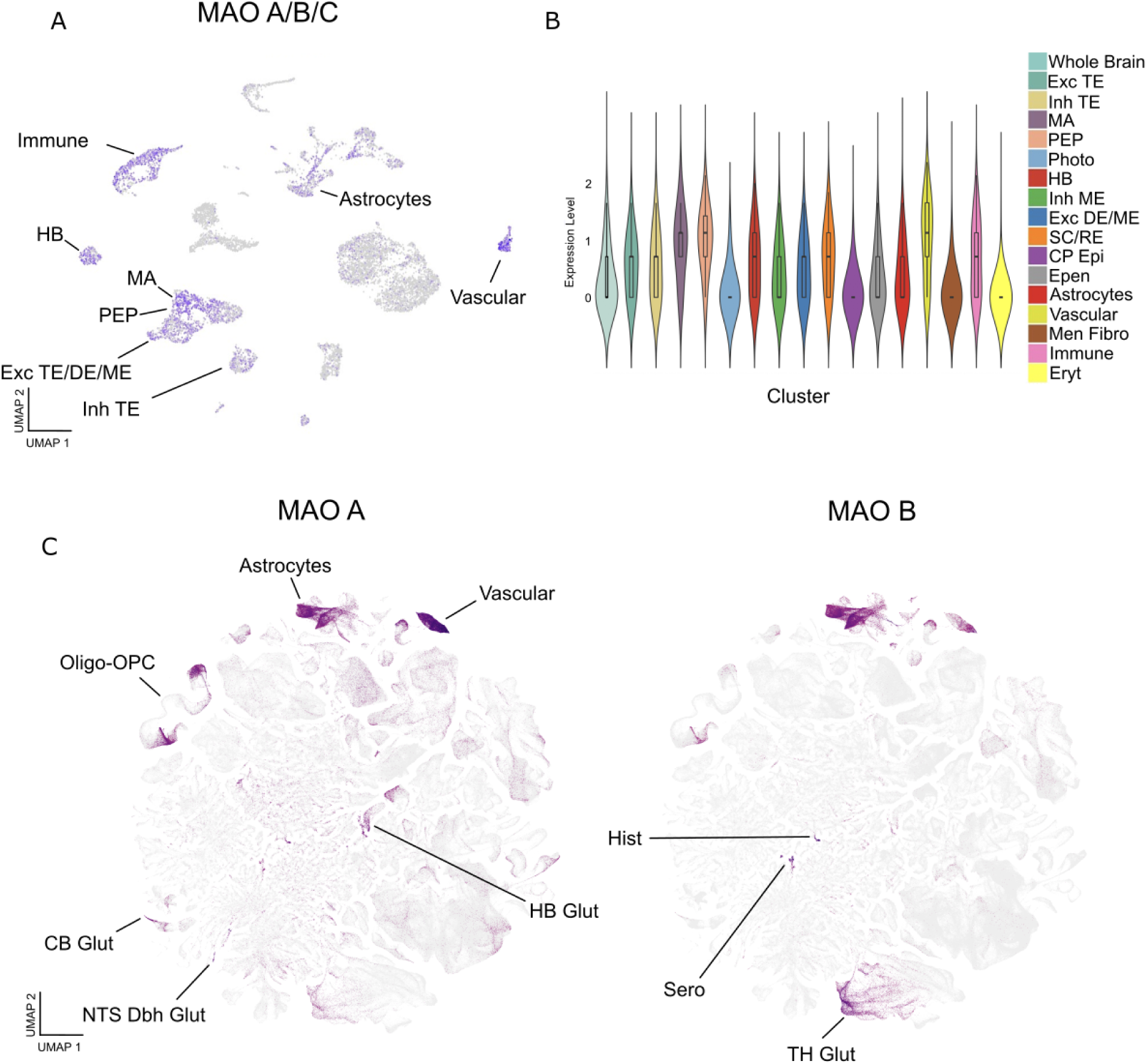
Expression of MAO A and MAO B in the mouse brain, and MAO A/B/C in the lamprey brain. (A) UMAP plot depicting the expression of lamprey MAO A/B/C in immune cells, habenular neurons (HB), monoaminergic neurons (MA), peptidergic neurons (PEP), excitatory neurons from the telencephalon, diencephalon and mesencephalon (Exc TE/DE/ME), inhibitory neurons from the telencephalon (Inh TE), astrocytes and vascular cells. (B) Violin plot depicting the expression levels of lamprey MAO A/B/C in specific cell clusters. Photoreceptors (Photo), spinal cord (SC), rhomboencephalon (RE), choroid plexus epithelium (CP Epi), ependymal cells (Epen), meningeal fibroblasts (Men Fibro), erythrocytes (Eryt). (C) Left: UMAP plot depicting the expression of mouse MAO A in vascular cells, astrocytes, oligodendrocytes and oligodendrocyte precursor cells (OPC), cerebellum glutamatergic neurons (CB Glut), dopamine-β-hydroxylase expressing glutamatergic neurons from the nucleus of the solitary tract (NTS Dbh Glut), habenular neurons (HB Glut). Right: UMAP plot depicting the expression of mouse MAO B in histaminergic neurons (Hist), serotonergic neurons (Sero), and glutamatergic neurons from the thalamus (TH Glut). The graphs do not include expression values lower than 7.6 Log2(CPM+1). Graphs in (C) were obtained from The Allen Brain Cell Atlas (https://portal.brain-map.org/atlases-and-data/bkp/abc-atlas).

Together, this data shows that while MAO A and MAO B are co-expressed in some cell types in the brain, they are also differentially enriched in others, as previously reported (Tong et al., 2013); (Levitt et al., 1982); (Konradi et al., 1988); (Shih et al., 1999)). For instance, while both show high abundance in vascular cells, astrocytes, and histaminergic neurons, MAO A shows higher expression in glutamatergic neurons from the cerebellum and brainstem neurons expressing noradrenergic markers. MAO B shows more relative enrichment in serotonergic neurons and glutamatergic neurons from the thalamus. This is in agreement with previous reports detecting MAO A mainly in catecholaminergic neurons and MAO B in serotonergic neurons and glia ((Levitt et al., 1982); (Konradi et al., 1988); (Shih et al., 1999)). Lamprey MAO A/B/C expression shows some similarities when compared to house mouse MAOs. This is especially reflected by its high expression in vascular cells and monoaminergic neurons. The role of MAO A/B/C in these cell types is currently unknown. Altogether, this suggests that, consistent with previous results, a gene duplication event in the tetrapod ancestor resulted in two MAO proteins with specialized but, in some cases, overlapping brain functions. Additionally, comparison with sequencing data from the sea lamprey brain strongly suggests a role for the vertebrate ancestor MAO protein in vascular cells and monoaminergic neurons.

### Conclusions

Our study provides new insights into the evolutionary relationships of the MAO gene family, revealing the existence of a new enzyme member (MAO C). Thus, most jawed vertebrates possess a repertoire of two MAO genes, MAO A and MAO B in tetrapods and MAO A/B and MAO C in non-tetrapod jawed vertebrates. Jawless vertebrates possess the ancestral condition of a single copy gene, MAO A/B/C. Using the Indo-pacific tarpon (*Megalops cyprinoides*) as a representative model system, we demonstrated that MAO A/B and MAO C localize to the outer mitochondrial membrane and exhibit enzymatic and structural properties comparable to human proteins. Notably, while the catalytic efficiency of MAO C is comparable to human MAOs (with a substrate specificity pattern more similar to human MAO A), MAO A/B features a remarkably higher enzymatic rate. This observation raised the hypothesis that this MAO arrangement in non-tetrapod jawed vertebrates was favored by increased oxygen concentrations in oceans in prehistoric ages, enabling these animals to respond rapidly to neuroactive amines in an increasingly complex environment. Extending these analyses to MAO A/B and MAO C family members in other non-tetrapod jawed organisms will allow us to probe if this MAO enzymatic machinery is a shared property and shed light on evolutionary mechanisms. Finally, our results strongly suggests a role for the vertebrate ancestor MAO protein in vascular cells and monoaminergic neurons.

## Materials and Methods

### Sequences and phylogenetic analyses

We obtained monoamine oxidase sequences from the National Center for Biotechnology Information (NCBI) (Sharma et al., 2018). For protein sequences, we used the human (*Homo sapiens*) MAO A and MAO B sequences as the reference for protein-BLAST (blastp) (Altschul et al., 1990) against the non-redundant database (nr) with default parameters. To further enrich our searches, we used sequences retrieved from different phylogenetic groups as a query for Blastp. For nucleotide sequences, we retrieved the corresponding DNA coding sequence from the accession numbers of the protein sequences mentioned above.

We implemented two types of analyses that involved different sampling strategies. The first aimed to understand the evolutionary history of the MAO gene family in vertebrates. Thus, our taxonomic sampling included protein sequences from representative species of all main groups of vertebrates (mammals, reptiles, birds, amphibians, lungfish, coelacanth, bony fish, cartilaginous fish, and jawless fish). The second analysis aimed to resolve better the evolution of MAO A and MAO B gene lineages in tetrapods. Thus, our taxonomic sampling included nucleotide sequences from representative species of all main groups of tetrapods (mammals, reptiles, birds, and amphibians). Accession numbers and details about the taxonomic sampling are available in Supplementary Table S1.

We used the software MAFFT v.7 (Katoh and Standley, 2013) to align sequences, allowing the program to choose the alignment strategies (L-INS-i for amino acids and FFT-NS-i for nucleotides). We estimated phylogenetic relationships using the maximum likelihood (ML) approach implemented in IQ-Tree v1.6.12 (Nguyen et al., 2015). We used the model finder tool of IQ-Tree v.1.6.12 (Kalyaanamoorthy et al., 2017) to select the best-fitting substitution models, which selected JTT+I+G4 for the amino acid alignment and GTR+F+R5 for the nucleotide alignment. We assessed the node support using the approximate Bayes test (Anisimova and Gascuel, 2006; Guindon, 2010) and ultrafast bootstrap approximation with 1000 pseudoreplicates (Hoang et al., 2018; Minh et al., 2013). We carried out 15 independent runs to explore the tree space, changing the strength value of the perturbation (-pers) parameter (0.3, 0.5, and 0.7). In all runs, the number of unsuccessful iterations to stop parameter (-nstop) value was changed from 100 (default value) to 500. The tree with the highest likelihood score was chosen. Sequences from human (*Homo sapiens*) and house mouse (*Mus musculus*) lysine-specific demethylase 1 (LSD1) and lysine-specific demethylase 2 (LSD2) were used as outgroup for the vertebrate phylogenetic tree, and coelacanth (*Latimeria chalumnae*) and West African lungfish MAO A/B sequences were used for the tetrapod tree.

### Assessment of conserved synteny

We examined genes found up- and downstream of MAO genes in species representative of gnathostomes. Synteny assessments were conducted using Genomicus v100.01 (Nguyen et al., 2018), the NCBI database (Sharma et al., 2018), and Ensembl v.108 (Cunningham et al., 2021). Our analyses included Human (*Homo sapiens*), Chicken (*Gallus gallus*), Green anole (*Anolis carolinensis*), Western clawed frog (*Xenopus tropicalis*), Zebrafish (*Danio rerio*), Spotted gar (*Lepisosteus oculatus*), Sterlet sturgeon (*Acipenser ruthenus*), Reedfish (*Erpetoichthys calabaricus*) and Whitespotted bamboo shark (*Chiloscyllium plagiosum*).

### Recombinant cDNA constructs

For expression in mammalian cells, codon-optimized mammalian expression constructs encoding full-length MAO A/B (amino acids 1–522) or MAO C (amino acids 1–528) from the Indo-pacific tarpon (*Megalops cyprinoides*) cloned in-frame into the *Eco*RV site of the pCMV-3Tag-2a vector, were acquired from GenScript (Piscataway, NJ). This configuration adds three successive Myc epitope tags encoded by the vector at the N-terminus of the expressed proteins. For the expression and purification of recombinant proteins in *Pichia pastoris*, DNA fragments encoding IPT MAO A/B and IPT MAO C were inserted into a pPIC3.5K vector (ThermoFisher) with the addition of a 6xHis tag sequence at the N-terminus. The pPIC3.5K plasmids containing the cloned MAO sequences were linearized with *Sal*I and transformation in *Pichia pastoris* was carried out by electroporation using KM71 strain (ThermoFisher).

### Cell Culture, Cell Transfection, and Fluorescence Microscopy

H4 human neuroglioma cells and HeLa human cervix adenocarcinoma cells were obtained from the American Type Culture Collection (Manassas, VA). Cells were cultured and transfected using standard methods as we have described previously (Opazo et al., 2023). Briefly, cells were maintained in Dulbecco’s modified Eagle’s medium (DMEM) supplemented with 10% heat-inactivated fetal bovine serum, 100 U/ml penicillin, 100 μg/ml streptomycin (ThermoFisher), and 5 μg/ml plasmocin (InvivoGen, San Diego, CA), and cultured in a humidified incubator with 5% CO_2_ at 37 °C. Transient transfections were performed in cells grown on top of glass coverslips on 24-well plates or grown on 6-well plates. Transfections were performed with Lipofectamine 2000 according to the manufacturer’s instructions (ThermoFisher) when cells reached ∼60% of confluence. For fluorescence microscopy and mitochondrial fluorescent staining, we used a protocol that we have also described previously (Opazo et al., 2023). Briefly, cells grown on glass coverslips, after 16-h of transfection, were treated for 30 min with MitoTracker Orange CMTMRos (ThermoFisher) according to the manufacturer’s instructions. After washing with phosphate-buffered saline (PBS) supplemented with 0.1 mM CaCl_2_ and 1 mM MgCl_2_ (PBS-CM), cells were fixed in 4% paraformaldehyde for 30 min at room temperature, permeabilized with 0.2% Triton X-100 in PBS-CM for 15 min at room temperature, and incubated with blocking solution (0.2% gelatin in PBS-CM) for 10 min at room temperature. Cells were incubated with a monoclonal antibody to the c-Myc epitope (1:200 dilution; clone 9E10; Covance, Princeton, NJ) for 30 min at 37 °C in a humidified chamber. After washing with PBS, cells were incubated with Alexa-Fluor-488-conjugated donkey anti-mouse IgG (1:1000 dilution) (ThermoFisher). After additional washing with PBS, coverslips were mounted onto glass slides with Fluoromount-G mounting medium (ThermoFisher). Fluorescence microscopy images were acquired also as we have described before (Opazo et al., 2023), with an AxioObserver.D1 microscope equipped with a PlanApo 63x oil immersion objective (NA 1.4) and an AxioCam MRm digital camera (Carl Zeiss). Quantitative analysis of the colocalization of fluorescence signals was also performed as previously described by us (Opazo et al., 2023), for which we obtained the Pearson’s correlation coefficient of pairwise comparisons from images acquired under identical settings, avoiding signal saturation and correcting for background, crosstalk, and noise signals on each set of images, using the plugin JACoP (Bolte and Cordelières 2006) implemented in the software ImageJ (version 1.47 h; Schneider et al. 2012). We prepared figures using 12-bit images processed with the software ImageJ and Adobe Photoshop CS3 (Adobe Systems, Mountain View, CA).

### Immunoblot analysis

Cells grown on 6-well plates, either non-transfected or after 16-h of transfection, were washed in ice-cold PBS and lysed at 4 °C in lysis buffer (50 mM Tris-HCl, 150 mM NaCl, 1 mM EDTA, 1% (v/v) Triton X-100, pH 7.4) supplemented with a cocktail of protease inhibitors (Sigma-Aldrich; cat # P8340). Lysates were cleared by centrifugation at 16,000 × *g* for 20 min at 4 °C. Samples with an equivalent amount of proteins were incubated at 65 °C for 5 min with Laemmli sample buffer (50 mM Tris-HCl, 2% SDS, 10% glycerol, 5% β-mercaptoethanol, 0.05% bromophenol blue, pH 6.8), and then processed by SDS-PAGE. Proteins were electrotransferred onto nitrocellulose membranes, followed by incubation in PBS containing 5% (w/v) nonfat dry milk (PBS-M) for 1-h at room temperature. Nitrocellulose membranes were incubated overnight at 4 °C with primary antibodies diluted in PBS-M, followed by six washes at room temperature in PBS containing 0.02% Tween-20 (PBS-T). Incubation with secondary antibodies also diluted in PBS-M was performed for 1-h at room temperature, followed by six washes in PBS-T at room temperature. Immunoreactive bands were detected by chemiluminescence using SuperSignal West Pico (Thermo Fisher Scientific). The primary antibodies used were mouse monoclonal anti-Myc epitope (Covance, clone 9E10; 1:1000 dilution) and mouse monoclonal anti-β-ACTIN used as an internal loading control (ThermoFisher, clone BA3R; 1:5000 dilution). The secondary antibody was horseradish peroxidase (HRP)-conjugated goat anti-mouse IgG (Jackson Immunoresearch, West Grove, PA, USA; cat # 715-035-150; 1:10000 dilution).

### Over-expression in *Pichia pastoris* and purification of IPT-MAO A/B and IPT-MAO C

His+ transformants were selected on minimal media plates without histidine, followed by a random selection of eight colonies for small-scale expression analysis. The transformants yielding the highest protein expression level were chosen for scale-up cultures. IPT MAO A/B and IPT MAO C were expressed and purified following a modified procedure of published protocols (Edmondson, 2023). A sample of the selected clone was smeared on a MD-Agar plate and incubated for two days at 30 °C. A single colony was inoculated in 5 ml YPD and incubated overnight at 30 °C (250 rpm shaking). A total of 15 ml of pre-culture was inoculated into flasks, each containing 330 ml of BMG medium (100 mM potassium phosphate pH 6.0, 4 x 10-5% w/v biotin, 1.34% w/v Yeast Nitrogen Base, 1% v/v glycerol), and left to grow at 30 °C with shaking at 200 rpm for three days. Cells were collected by centrifugation at 5,000 x g for 10 min and resuspended in 165 ml of BMM medium (100 mM potassium phosphate pH 6.0, 4 x 10-5% w/v biotin, 1.34% w/v Yeast Nitrogen Base, 0.5% v/v methanol) for each flask to induce protein expression. A supplement of 825 μl methanol for each flask was added every 12 h. After induction, cells were collected by centrifugation at 5,000 x g for 15 min, flash-frozen in liquid nitrogen and stored -80 °C.

For purification, 20 g of cells were resuspended in 100 ml of breaking buffer (50 mM sodium phosphate pH 7.2, 5 % w/v glycerol, 1 mM PSMF, 30 μM DTT, 1 mM EDTA) with an equal volume of silica-zirconia beads (0.5 mm in diameter) and then disrupted in a Beadbeater (Hamilton Beach Blender 908). The cell lysate was separated from beads through filtration with a Miracloth (Calbiochem) layer followed by low-speed centrifugation (1500 x g for 10 min at 4 °C). The supernatant was centrifuged at high speed (70000 x g for 45 min at 4 °C) in order to separate the membrane fraction. The pellet was resuspended to a final concentration of 15 mg/ml in 50 mM sodium phosphate pH 7.8, 300 mM sodium chloride, 20 mM imidazole, 20 % w/v glycerol, 1 mM PMSF, 30 μM DTT. The protein concentration was determined using the Biuret method. Detergent extraction was carried out by adding 1 % w/v FOS-choline12 (Anatrace) and the mixture was stirred at 4 °C in the dark for 1 h. After centrifugation (70000 x g for 30 min) the extract was loaded onto a 5 ml Nickel column (Cytiva) equilibrated with breaking buffer using an AktaPure system (Cytiva). After washing with 10 column volumes of the same buffer, the enzyme was eluted with 50 mM sodium phosphate buffer pH 7.8, 300 mM sodium chloride, 300 mM imidazole, 20 % glycerol w/v, 1 mM PMSF, 30 μM DTT. Elution was followed by monitoring absorbance at 280 nm and 456 nm, the latter being the absorption wavelength of the MAO flavin cofactor. The enzyme fractions corresponding to the elution peak were pooled and concentrated with Amicon Ultra 30K centrifugal filters (Millipore). The excess of imidazole was removed with a 5 ml HiTrap Desalting column (Cytiva) equilibrated with 50 mM sodium phosphate buffer pH 7.8, 300 mM sodium chloride, 20 % w/v glycerol. The purity of the protein was determined by SDS-PAGE analysis and inspection of the UV-Vis spectrum. The final concentration of the pure protein was obtained spectrophotometrically with a NanoDrop ND-14081000 (ThermoFisher Scientific Inc.) from the absorbance of the enzyme-bound FAD coenzyme at 456 nm (ε456 = 12000 M-1 cm-1). Recombinant human MAO A and MAO B were over-expressed and purified using published protocols (Edmondson, 2023).

### Enzymatic and inhibition assays

The enzymatic activity of both IPT MAOs was measured by a spectrophotometric assay in the presence of varying concentrations of substrates using a Cary 100 UV/Vis spectrophotometer (Agilent Technologies, CA, USA) (Reis and Binda, 2023). All experiments were performed at 25 °C in 50 mM HEPES/NaOH pH 7.5 containing 0.5 % v/v reduced Triton X-100. The rate of substrate oxidation was monitored spectrophotometrically at 515 nm (ε515 = 26 mM-1 cm-1) using the horseradish peroxidase (HRP) coupled assay that monitors the production of hydrogen peroxide. The assay mixture contained 0.10 mM 4-aminoantipyrine, 1.0 mM 3,5-dichloro-2-hydroxybenzenesulfonic acid, and 0.013 mg ml-1 HRP. The reaction was started by adding the enzyme at a final concentration ranging from 0.1 μM and 1 μM depending on the substrate. Steady-state parameters for IPT MAOs with dopamine and serotonin were determined by monitoring oxygen consumption using an Oxygraph plus system (Hansatech Instruments Ltd.) because these molecules are known to interfere with the spectrophotometric assay by binding to HRP. Measurements were performed in 50 mM HEPES/NaOH buffer pH 7.5, 0.5 v/v reduced Triton X-100, using 0.5 μM enzyme while varying the substrate concentration between 0.5 and 15 mM. The reaction chamber had a total volume of 1.0 ml, and the sample was constantly stirred (60 rpm). The reactions were initiated by adding the substrate and followed by monitoring the initial linear decrease in oxygen concentration. All data were analyzed by non-linear regression curve with GraphPad Prism 8.0 software (La Jolla, CA, USA) using Michaelis-Menten curve model. Irreversible inhibition of IPT MAOs, using the well-known inhibitors clorgyline (human MAO A selective) and deprenyl (human MAO B selective) was tested by UV-Vis spectral measurement using an Agilent 8453 UV/Vis diode-array spectrophotometer. The experiments were performed in a quartz cuvette containing a solution of 10 μM enzyme in 50 mM potassium phosphate pH 7.8, 300 mM sodium chloride, 20% w/v glycerol, 0.05% w/v FOS-Choline-12 and following the absorbance spectrum after addition of a 10-fold molar excess of irreversible inhibitor. The reaction was monitored at different times until no further modification was detected. The final spectra and time course of absorbance changes observed with each inhibitor were plotted using GraphPad 8.0 software.

### Single-cell RNAseq analysis

The whole brain of the adult lamprey dataset was obtained from the Lamprey brain atlas developed by the Kaessmann Lab (https://lampreybrain.kaessmannlab.org/). We downloaded the dataset as an RDS file, which was analyzed using Seurat v.5.0.3 (Hao et al., 2024). Normalization was performed using the SCTransform method (Hafemeister and Satija, 2019) to select the top 3,000 highly variable genes. Subsequently, we reduced the dimensionality of the data with the Principal Component Analysis (PCA), and we utilized the first 30 principal components to build the shared nearest neighbor that was clustered using the Louvain method. To visualize the resulting clusters in two-dimensions, we computed the Uniform Manifold Approximation and Projection (UMAP). We maintained the original identities of the clusters that were manually annotated and reported in (Lamanna et al., 2023). For the mouse brain dataset, we constructed the figures using The Allen Brain Cell Atlas (https://portal.brain-map.org/atlases-and-data/bkp/abc-atlas), which allows access to data from (Yao et al., 2023).

## Acknowledgments

This work was supported by Fondo Nacional de Desarrollo Científico y Tecnológico from Chile, FONDECYT 1210471 to JCO, and FONDECYT 1211481 to GAM. The PhD fellowship of LB (38th cycle) was funded by PNRR (Missione 4, componente 1, Investimento 3.4), D.M. n. 351 April 9th 2022. We thank Charlotte Luchsinger and Gonzalo Astroza for their technical assistance. This article is dedicated to the memory of Roberto Araya Ariztía.

---

**Supplementary Figure S1.**
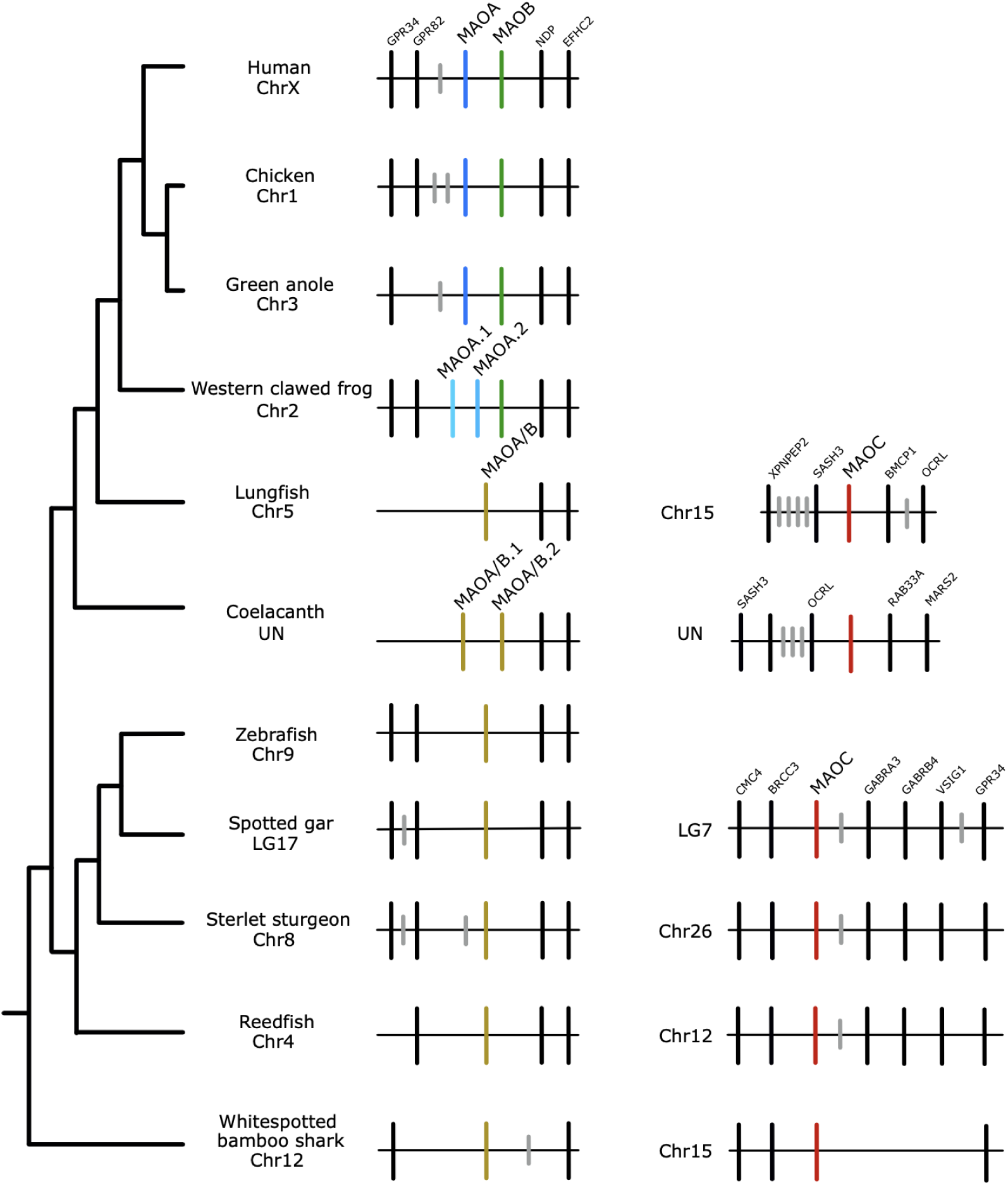
Patterns of conserved synteny in the chromosomal regions harboring the monoamine oxidase genes of jawed vertebrates. Gray lines represent genes that do not contribute to conserved synteny. UN: Unplaced scaffold.

**Supplementary Figure S2.**
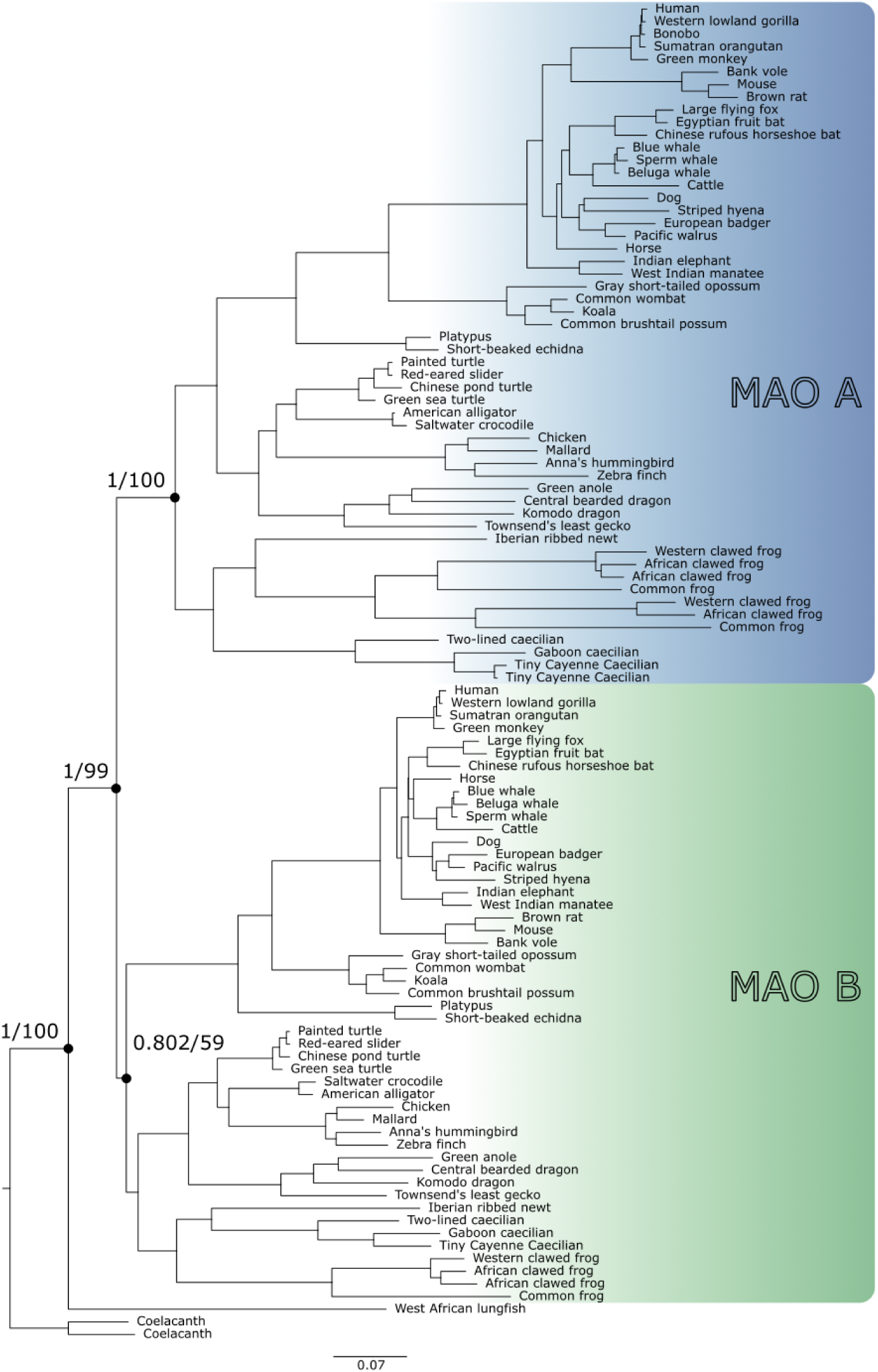
Maximum-likelihood tree showing relationships among monoamine oxidases of tetrapods. Numbers on the nodes correspond to support from the abayes and ultrafast bootstrap values. The scale denotes substitutions per site, and shading represents gene lineages. MAO A/B nucleotide sequences from the West African lungfish (*Protopterus annectens*) and coelacanth (*Latimeria chalumnae*) were used as outgroups.

**Supplementary Figure S3.**
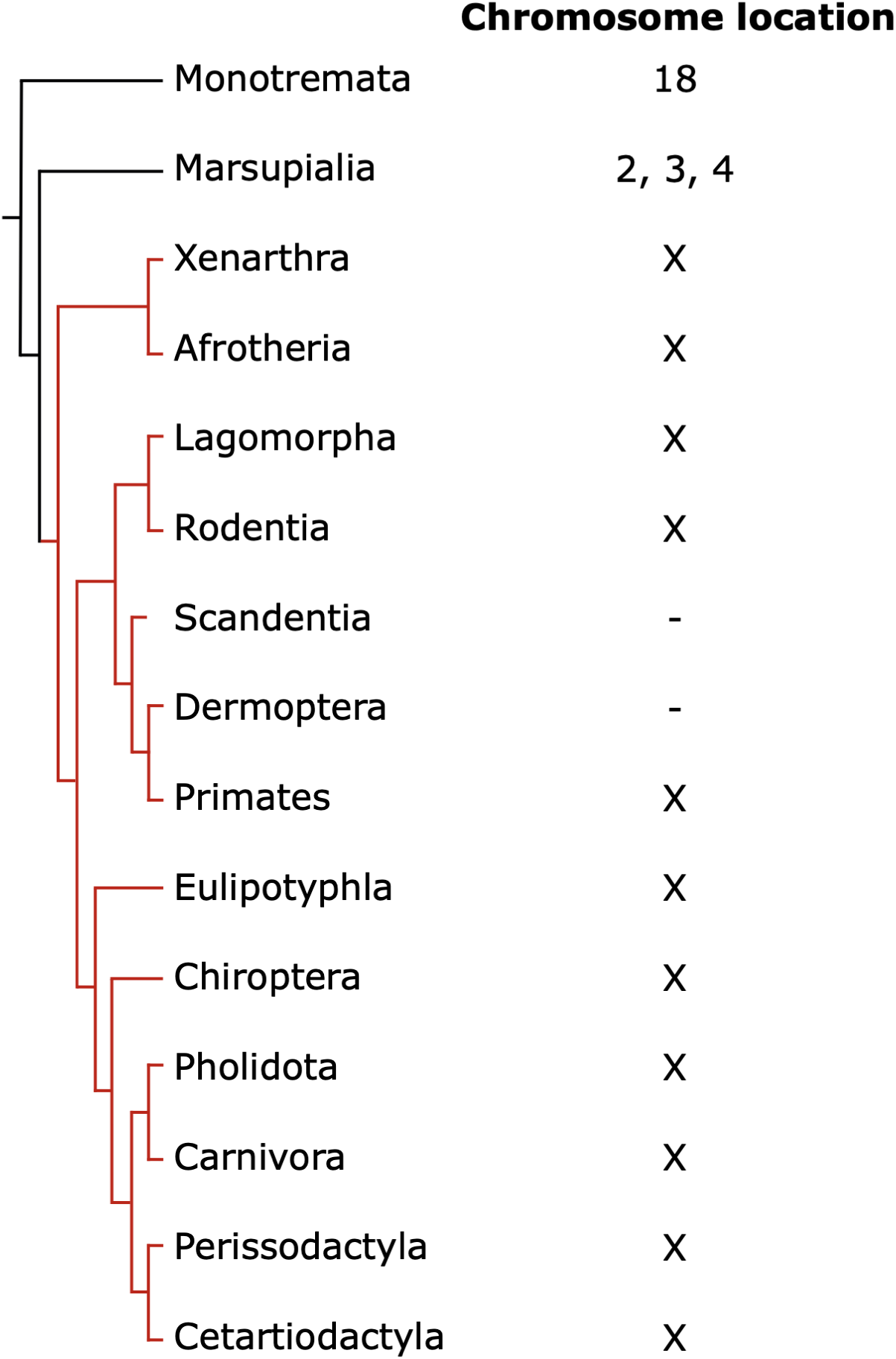
Chromosomal location of MAO A and MAO B genes in mammals. As is shown, the translocation of MAO genes from an autosome to chromosome X occurred in the last common ancestor of placental mammals between 160 and 99 million years ago (Kumar et al., 2022). The tree topology was obtained from the literature (Esselstyn et al., 2017). Placental mammals are indicated with red branches.

**Supplementary Figure S4.**
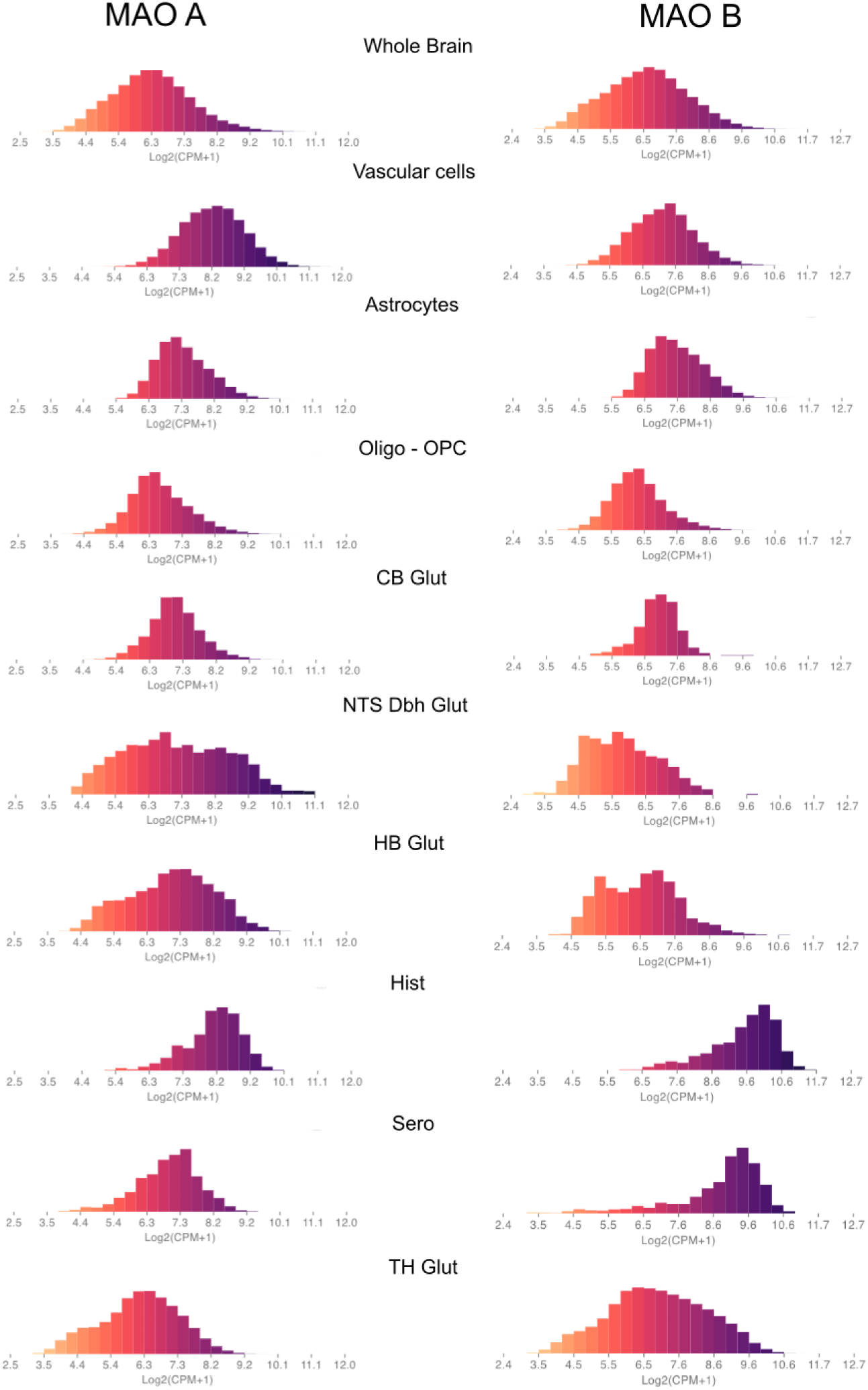
Expression of MAO A and MAO B in the mouse brain. Distribution of expression values of MAO A and MAO B transcripts in different mouse brain cell types. Oligo: oligodendrocytes, OPC: Oligodendrocyte precursor cells, CB: Cerebellum, Glut: Glutamatergic, NTS: Nucleus of the solitary tract, Dbh: Dopamine-β-hydroxylase, HB: Habenula, Hist: Histaminergic, Sero: Serotonergic, TH: Thalamus. These graphs were obtained from The Allen Brain Cell Atlas (https://portal.brain-map.org/atlases-and-data/bkp/abc-atlas).

## References

Abramson J, Adler J, Dunger J, Evans R, Green T, Pritzel A, Ronneberger O, Willmore L, Ballard AJ, Bambrick J, Bodenstein SW, Evans DA, Hung C-C, O’Neill M, Reiman D, Tunyasuvunakool K, Wu Z, Žemgulytė A, Arvaniti E, Beattie C, Bertolli O, Bridgland A, Cherepanov A, Congreve M, Cowen-Rivers AI, Cowie A, Figurnov M, Fuchs FB, Gladman H, Jain R, Khan YA, Low CMR, Perlin K, Potapenko A, Savy P, Singh S, Stecula A, Thillaisundaram A, Tong C, Yakneen S, Zhong ED, Zielinski M, Žídek A, Bapst V, Kohli P, Jaderberg M, Hassabis D, Jumper JM. 2024. Accurate structure prediction of biomolecular interactions with AlphaFold 3. Nature 630:493–500.

Aldeco M, Arslan BK, Edmondson DE. 2011. Catalytic and inhibitor binding properties of zebrafish monoamine oxidase (zMAO): comparisons with human MAO A and MAO B. Comp Biochem Physiol B Biochem Mol Biol 159:78–83.

Altenhoff AM, Warwick Vesztrocy A, Bernard C, Train C-M, Nicheperovich A, Prieto Baños S, Julca I, Moi D, Nevers Y, Majidian S, Dessimoz C, Glover NM. 2024. OMA orthology in 2024: improved prokaryote coverage, ancestral and extant GO enrichment, a revamped synteny viewer and more in the OMA Ecosystem. Nucleic Acids Res 52:D513–D521.

Altschul SF, Gish W, Miller W, Myers EW, Lipman DJ. 1990. Basic local alignment search tool. J Mol Biol 215:403–410.

Anctil M. 2009. Chemical transmission in the sea anemone Nematostella vectensis: A genomic perspective. Comp Biochem Physiol Part D Genomics Proteomics 4:268–289.

Anichtchik O, Sallinen V, Peitsaro N, Panula P. 2006. Distinct structure and activity of monoamine oxidase in the brain of zebrafish (Danio rerio). J Comp Neurol 498:593–610.

Anisimova M, Gascuel O. 2006. Approximate likelihood-ratio test for branches: A fast, accurate, and powerful alternative. Syst Biol 55:539–552.

Bach AW, Lan NC, Johnson DL, Abell CW, Bembenek ME, Kwan SW, Seeburg PH, Shih JC. 1988. cDNA cloning of human liver monoamine oxidase A and B: molecular basis of differences in enzymatic properties. Proc Natl Acad Sci U S A 85:4934–4938.

Chen S, Krinsky BH, Long M. 2013. New genes as drivers of phenotypic evolution. Nat Rev Genet 14:645–660.

Cunningham F, Allen JE, Allen J, Alvarez-Jarreta J, Amode MR, Armean IM, Austine-Orimoloye O, Azov AG, Barnes I, Bennett R, Berry A, Bhai J, Bignell A, Billis K, Boddu S, Brooks L, Charkhchi M, Cummins C, Da Rin Fioretto L, Davidson C, Dodiya K, Donaldson S, El Houdaigui B, El Naboulsi T, Fatima R, Giron CG, Genez T, Martinez JG, Guijarro-Clarke C, Gymer A, Hardy M, Hollis Z, Hourlier T, Hunt T, Juettemann T, Kaikala V, Kay M, Lavidas I, Le T, Lemos D, Marugán JC, Mohanan S, Mushtaq A, Naven M, Ogeh DN, Parker A, Parton A, Perry M, Piližota I, Prosovetskaia I, Sakthivel MP, Salam AIA, Schmitt BM, Schuilenburg H, Sheppard D, Pérez-Silva JG, Stark W, Steed E, Sutinen K, Sukumaran R, Sumathipala D, Suner M-M, Szpak M, Thormann A, Tricomi FF, Urbina-Gómez D, Veidenberg A, Walsh TA, Walts B, Willhoft N, Winterbottom A, Wass E, Chakiachvili M, Flint B, Frankish A, Giorgetti S, Haggerty L, Hunt SE, IIsley GR, Loveland JE, Martin FJ, Moore B, Mudge JM, Muffato M, Perry E, Ruffier M, Tate J, Thybert D, Trevanion SJ, Dyer S, Harrison PW, Howe KL, Yates AD, Zerbino DR, Flicek P. 2021. Ensembl 2022. Nucleic Acids Res 50:D988–D995.

De Colibus L, Li M, Binda C, Lustig A, Edmondson DE, Mattevi A. 2005. Three-dimensional structure of human monoamine oxidase A (MAO A): relation to the structures of rat MAO A and human MAO B. Proc Natl Acad Sci U S A 102:12684–12689.

Demuth JP, De Bie T, Stajich JE, Cristianini N, Hahn MW. 2006. The evolution of mammalian gene families. PLoS One 1:e85.

Edmondson DE. 2023. Purification of Recombinant Eukaryotic MAO A and MAO B Utilizing the Pichia pastoris Expression System. Methods Mol Biol 2558:11–22.

Edmondson DE, Binda C. 2018. Monoamine Oxidases. Subcell Biochem 87:117–139.

Edmondson DE, Binda C, Wang J, Upadhyay AK, Mattevi A. 2009. Molecular and mechanistic properties of the membrane-bound mitochondrial monoamine oxidases. Biochemistry 48:4220–4230.

Emms DM, Kelly S. 2019. OrthoFinder: phylogenetic orthology inference for comparative genomics. Genome Biol 20:238.

Esselstyn JA, Oliveros CH, Swanson MT, Faircloth BC. 2017. Investigating Difficult Nodes in the Placental Mammal Tree with Expanded Taxon Sampling and Thousands of Ultraconserved Elements. Genome Biol Evol 9:2308–2321.

Goodman M, Czelusniak J, Moore GW, Romero-Herrera AE, Matsuda G. 1979. Fitting the gene lineage into its species lineage, a parsimony strategy illustrated by cladograms constructed from globin sequences. Syst Zool 28:132.

Goulty M, Botton-Amiot G, Rosato E, Sprecher SG, Feuda R. 2023. The monoaminergic system is a bilaterian innovation. Nat Commun 14:3284.

Guindon S. 2010. Bayesian estimation of divergence times from large sequence alignments. Mol Biol Evol 27:1768–1781.

Hafemeister C, Satija R. 2019. Normalization and variance stabilization of single-cell RNA-seq data using regularized negative binomial regression. Genome Biol 20:296.

Hao Y, Stuart T, Kowalski MH, Choudhary S, Hoffman P, Hartman A, Srivastava A, Molla G, Madad S, Fernandez-Granda C, Satija R. 2024. Dictionary learning for integrative, multimodal and scalable single-cell analysis. Nat Biotechnol 42:293–304.

Himmel NJ, Cox DN. 2020. Transient receptor potential channels: current perspectives on evolution, structure, function and nomenclature. Proc Biol Sci 287:20201309.

Himmel NJ, Gray TR, Cox DN. 2020. Phylogenetics Identifies Two Eumetazoan TRPM Clades and an Eighth TRP Family, TRP Soromelastatin (TRPS). Mol Biol Evol 37:2034–2044.

Hoang DT, Chernomor O, von Haeseler A, Minh BQ, Vinh LS. 2018. UFBoot2: Improving the Ultrafast Bootstrap Approximation. Mol Biol Evol 35:518–522.

Iacovino LG, Manzella N, Resta J, Vanoni MA, Rotilio L, Pisani L, Edmondson DE, Parini A, Mattevi A, Mialet-Perez J, Binda C. 2020. Rational Redesign of Monoamine Oxidase A into a Dehydrogenase to Probe ROS in Cardiac Aging. ACS Chem Biol 15:1795–1800.

Iagodina OV, Basova IN. 2013. [Liver monoamine oxidase activity of the lamprey Lampetra fluviatilis. the substrate-inhibitory specificity]. Zh Evol Biokhim Fiziol 49:39–43.

Jones DN, Raghanti MA. 2021. The role of monoamine oxidase enzymes in the pathophysiology of neurological disorders. J Chem Neuroanat 114:101957.

Kalyaanamoorthy S, Minh BQ, Wong TKF, von Haeseler A, Jermiin LS. 2017. ModelFinder: fast model selection for accurate phylogenetic estimates. Nat Methods 14:587–589.

Katoh K, Standley DM. 2013. MAFFT multiple sequence alignment software version 7: improvements in performance and usability. Mol Biol Evol 30:772–780.

Kearney EB, Salach JI, Walker WH, Seng RL, Kenney W, Zeszotek E, Singer TP. 1971. The covalently-bound flavin of hepatic monoamine oxidase. 1. Isolation and sequence of a flavin peptide and evidence for binding at the 8alpha position. Eur J Biochem 24:321–327.

Kobayashi S, Takahara K, Kamijo K. 1981. Monoamine oxidase in frog liver and brain. Comp Biochem Physiol C 69:179–183.

Kokel D, Bryan J, Laggner C, White R, Cheung CYJ, Mateus R, Healey D, Kim S, Werdich AA, Haggarty SJ, Macrae CA, Shoichet B, Peterson RT. 2010. Rapid behavior-based identification of neuroactive small molecules in the zebrafish. Nat Chem Biol 6:231–237.

Konradi C, Svoma E, Jellinger K, Riederer P, Denney R, Thibault J. 1988. Topographic immunocytochemical mapping of monoamine oxidase-A, monoamine oxidase-B and tyrosine hydroxylase in human post mortem brain stem. Neuroscience 26:791–802.

Kuderna LFK, Gao H, Janiak MC, Kuhlwilm M, Orkin JD, Bataillon T, Manu S, Valenzuela A, Bergman J, Rousselle M, Silva FE, Agueda L, Blanc J, Gut M, de Vries D, Goodhead I, Harris RA, Raveendran M, Jensen A, Chuma IS, Horvath JE, Hvilsom C, Juan D, Frandsen P, Schraiber JG, de Melo FR, Bertuol F, Byrne H, Sampaio I, Farias I, Valsecchi J, Messias M, da Silva MNF, Trivedi M, Rossi R, Hrbek T, Andriaholinirina N, Rabarivola CJ, Zaramody A, Jolly CJ, Phillips-Conroy J, Wilkerson G, Abee C, Simmons JH, Fernandez-Duque E, Kanthaswamy S, Shiferaw F, Wu D, Zhou L, Shao Y, Zhang G, Keyyu JD, Knauf S, Le MD, Lizano E, Merker S, Navarro A, Nadler T, Khor CC, Lee J, Tan P, Lim WK, Kitchener AC, Zinner D, Gut I, Melin AD, Guschanski K, Schierup MH, Beck RMD, Umapathy G, Roos C, Boubli JP, Rogers J, Farh KK-H, Marques Bonet T. 2023. A global catalog of whole-genome diversity from 233 primate species. Science 380:906–913.

Kumar S, Suleski M, Craig JM, Kasprowicz AE, Sanderford M, Li M, Stecher G, Hedges SB. 2022. TimeTree 5: An Expanded Resource for Species Divergence Times. Mol Biol Evol 39. doi:10.1093/molbev/msac174

Kutchko KM, Siltberg-Liberles J. 2013. Metazoan innovation: from aromatic amino acids to extracellular signaling. Amino Acids 45:359–367.

Lamanna F, Hervas-Sotomayor F, Oel AP, Jandzik D, Sobrido-Cameán D, Santos-Durán GN, Martik ML, Stundl J, Green SA, Brüning T, Mößinger K, Schmidt J, Schneider C, Sepp M, Murat F, Smith JJ, Bronner ME, Rodicio MC, Barreiro-Iglesias A, Medeiros DM, Arendt D, Kaessmann H. 2023. A lamprey neural cell type atlas illuminates the origins of the vertebrate brain. Nat Ecol Evol 7:1714–1728.

Levitt P, Pintar JE, Breakefield XO. 1982. Immunocytochemical demonstration of monoamine oxidase B in brain astrocytes and serotonergic neurons. Proc Natl Acad Sci U S A 79:6385–6389.

Martin FJ, Amode MR, Aneja A, Austine-Orimoloye O, Azov AG, Barnes I, Becker A, Bennett R, Berry A, Bhai J, Bhurji SK, Bignell A, Boddu S, Branco Lins PR, Brooks L, Ramaraju SB, Charkhchi M, Cockburn A, Da Rin Fiorretto L, Davidson C, Dodiya K, Donaldson S, El Houdaigui B, El Naboulsi T, Fatima R, Giron CG, Genez T, Ghattaoraya GS, Martinez JG, Guijarro C, Hardy M, Hollis Z, Hourlier T, Hunt T, Kay M, Kaykala V, Le T, Lemos D, Marques-Coelho D, Marugán JC, Merino GA, Mirabueno LP, Mushtaq A, Hossain SN, Ogeh DN, Sakthivel MP, Parker A, Perry M, Piližota I, Prosovetskaia I, Pérez-Silva JG, Salam AIA, Saraiva-Agostinho N, Schuilenburg H, Sheppard D, Sinha S, Sipos B, Stark W, Steed E, Sukumaran R, Sumathipala D, Suner M-M, Surapaneni L, Sutinen K, Szpak M, Tricomi FF, Urbina-Gómez D, Veidenberg A, Walsh TA, Walts B, Wass E, Willhoft N, Allen J, Alvarez-Jarreta J, Chakiachvili M, Flint B, Giorgetti S, Haggerty L, Ilsley GR, Loveland JE, Moore B, Mudge JM, Tate J, Thybert D, Trevanion SJ, Winterbottom A, Frankish A, Hunt SE, Ruffier M, Cunningham F, Dyer S, Finn RD, Howe KL, Harrison PW, Yates AD, Flicek P. 2023. Ensembl 2023. Nucleic Acids Res 51:D933–D941.

McNicholas S, Potterton E, Wilson KS, Noble MEM. 2011. Presenting your structures: the CCP4mg molecular-graphics software. Acta Crystallogr D Biol Crystallogr 67:386–394.

Milczek EM, Binda C, Rovida S, Mattevi A, Edmondson DE. 2011. The “gating” residues Ile199 and Tyr326 in human monoamine oxidase B function in substrate and inhibitor recognition. FEBS J 278:4860–4869.

Minh BQ, Nguyen MAT, von Haeseler A. 2013. Ultrafast approximation for phylogenetic bootstrap. Mol Biol Evol 30:1188–1195.

Nguyen L-T, Schmidt HA, von Haeseler A, Minh BQ. 2015. IQ-TREE: a fast and effective stochastic algorithm for estimating maximum-likelihood phylogenies. Mol Biol Evol 32:268–274.

Nguyen NTT, Vincens P, Roest Crollius H, Louis A. 2018. Genomicus 2018: karyotype evolutionary trees and on-the-fly synteny computing. Nucleic Acids Res 46:D816–D822.

Nicotra A, Falasca L, Senatori O, Conti Devirgiliis L. 2002. Monoamine oxidase A and B activities in embryonic chick hepatocytes: differential regulation by retinoic acid. Cell Biochem Funct 20:87–94.

Opazo JC, Vandewege MW, Gutierrez J, Zavala K, Vargas-Chacoff L, Morera FJ, Mardones GA. 2021. Independent duplications of the Golgi phosphoprotein 3 oncogene in birds. Sci Rep 11:12483.

Opazo JC, Vandewege MW, Hoffmann FG, Zavala K, Meléndez C, Luchsinger C, Cavieres VA, Vargas-Chacoff L, Morera FJ, Burgos PV, Tapia-Rojas C, Mardones GA. 2023. How Many Sirtuin Genes Are Out There? Evolution of Sirtuin Genes in Vertebrates With a Description of a New Family Member. Mol Biol Evol 40. doi:10.1093/molbev/msad014

Pintar JE, Barbosa J, Francke U, Castiglione CM, Hawkins M Jr, Breakefield XO. 1981. Gene for monoamine oxidase type A assigned to the human X chromosome. J Neurosci 1:166–175.

Reis J, Binda C. 2023. The Peroxidase-Coupled Assay to Measure MAO Enzymatic Activity. Methods Mol Biol 2558:23–34.

Sayers EW, Bolton EE, Brister JR, Canese K, Chan J, Comeau DC, Connor R, Funk K, Kelly C, Kim S, Madej T, Marchler-Bauer A, Lanczycki C, Lathrop S, Lu Z, Thibaud-Nissen F, Murphy T, Phan L, Skripchenko Y, Tse T, Wang J, Williams R, Trawick BW, Pruitt KD, Sherry ST. 2022. Database resources of the national center for biotechnology information. Nucleic Acids Res 50:D20–D26.

Setini A, Pierucci F, Senatori O, Nicotra A. 2005. Molecular characterization of monoamine oxidase in zebrafish (Danio rerio). Comp Biochem Physiol B Biochem Mol Biol 140:153–161.

Sharma S, Ciufo S, Starchenko E, Darji D, Chlumsky L, Karsch-Mizrachi I, Schoch CL. 2018. The NCBI BioCollections Database. Database 2018. doi:10.1093/database/bay006

Shih JC, Chen K, Ridd MJ. 1999. Monoamine oxidase: from genes to behavior. Annu Rev Neurosci 22:197–217.

Tong J, Meyer JH, Furukawa Y, Boileau I, Chang L-J, Wilson AA, Houle S, Kish SJ. 2013. Distribution of monoamine oxidase proteins in human brain: implications for brain imaging studies. J Cereb Blood Flow Metab 33:863–871.

Vignieri S. 2023. Zoonomia. Science 380:356–357.

Wichmann IA, Zavala K, Hoffmann FG, Vandewege MW, Corvalán AH, Amigo JD, Owen GI, Opazo JC. 2016. Evolutionary history of the reprimo tumor suppressor gene family in vertebrates with a description of a new reprimo gene lineage. Gene 591:245–254.

Wu HF, Chen K, Shih JC. 1993. Site-directed mutagenesis of monoamine oxidase A and B: role of cysteines. Mol Pharmacol 43:888–893.

Yao Z, van Velthoven CTJ, Kunst M, Zhang M, McMillen D, Lee C, Jung W, Goldy J, Abdelhak A, Aitken M, Baker K, Baker P, Barkan E, Bertagnolli D, Bhandiwad A, Bielstein C, Bishwakarma P, Campos J, Carey D, Casper T, Chakka AB, Chakrabarty R, Chavan S, Chen M, Clark M, Close J, Crichton K, Daniel S, DiValentin P, Dolbeare T, Ellingwood L, Fiabane E, Fliss T, Gee J, Gerstenberger J, Glandon A, Gloe J, Gould J, Gray J, Guilford N, Guzman J, Hirschstein D, Ho W, Hooper M, Huang M, Hupp M, Jin K, Kroll M, Lathia K, Leon A, Li S, Long B, Madigan Z, Malloy J, Malone J, Maltzer Z, Martin N, McCue R, McGinty R, Mei N, Melchor J, Meyerdierks E, Mollenkopf T, Moonsman S, Nguyen TN, Otto S, Pham T, Rimorin C, Ruiz A, Sanchez R, Sawyer L, Shapovalova N, Shepard N, Slaughterbeck C, Sulc J, Tieu M, Torkelson A, Tung H, Valera Cuevas N, Vance S, Wadhwani K, Ward K, Levi B, Farrell C, Young R, Staats B, Wang M-QM, Thompson CL, Mufti S, Pagan CM, Kruse L, Dee N, Sunkin SM, Esposito L, Hawrylycz MJ, Waters J, Ng L, Smith K, Tasic B, Zhuang X, Zeng H. 2023. A high-resolution transcriptomic and spatial atlas of cell types in the whole mouse brain. Nature 624:317–332.

Yeung AWK, Georgieva MG, Atanasov AG, Tzvetkov NT. 2019. Monoamine Oxidases (MAOs) as Privileged Molecular Targets in Neuroscience: Research Literature Analysis. Front Mol Neurosci 12:143.

